# Large-scale RNAi screening uncovers new therapeutic targets in the human parasite *Schistosoma mansoni*

**DOI:** 10.1101/2020.02.05.935833

**Authors:** Jipeng Wang, Carlos Paz, Gilda Padalino, Avril Coghlan, Zhigang Lu, Irina Gradinaru, Julie N.R. Collins, Matthew Berriman, Karl F. Hoffmann, James J. Collins

## Abstract

Schistosomes kill 250,000 people every year and are responsible for serious morbidity in 240 million of the world’s poorest people. Despite their profound global impact, only a single drug (praziquantel) is available to treat schistosomiasis, highlighting the need to better understand schistosome biology to drive the development of a new generation of therapeutics. A major barrier to this goal is the paucity of large-scale datasets exploring schistosome gene function. Here, we describe the first large-scale RNA interference screen in adult *Schistosoma mansoni* examining the function of over 2000 genes representing approximately 20 percent of the protein coding genome. More than 250 genes were found to have phenotypes affecting neuromuscular function, tissue integrity, stem cell maintenance, and parasite survival. Leveraging these data, we bioinformatically prioritized several compounds with *in vitro* activity against parasites and validated p97, a component of the ubiquitin proteasome system, as a drug target in the worm. We further reveal a potentially druggable protein kinase-signaling module involving the TAO and STK25 kinases that are essential for maintaining the transcription of muscle-specific mRNAs. Importantly, loss of either of these kinases results in paralysis and death of schistosomes following surgical transplantation into a mammalian host. We anticipate this work will invigorate studies into the biology of these poorly studied organisms and expedite the development of new therapeutics to treat an important neglected tropical disease.

---

Genome sequences are available for the major species of medically-relevant schistosomes^1–3^; nevertheless, studies of gene function have been limited to relatively small numbers of genes^4,5^. To address this issue, we developed a platform for large-scale RNAi screening on adult schistosomes (**Fig. 1a**). To establish the efficacy of this platform to detect phenotypes in adult *S. mansoni*, we prioritized a list of 2,320 of the worm’s ~10,000 protein coding genes, including those encoding enzymes, cell-surface receptors, ion channels, and hypothetical proteins of unknown function (**Supplementary Table 1**). After filtering for genes expressed in adult schistosome somatic tissues using existing expression datasets^6^, we performed Polymerase Chain Reactions (PCR) from schistosome cDNA, generated dsRNAs, and performed RNAi by treating adult pairs of male and female worms with five dsRNA treatments over the course of a 30-day experiment (**Fig. 1b**). After filtering genes that either did not amplify during PCR steps, or failed to generate sufficient concentrations of dsRNA, a total of 2,216 genes were screened (**Supplementary Table 1**).

**Fig1.**
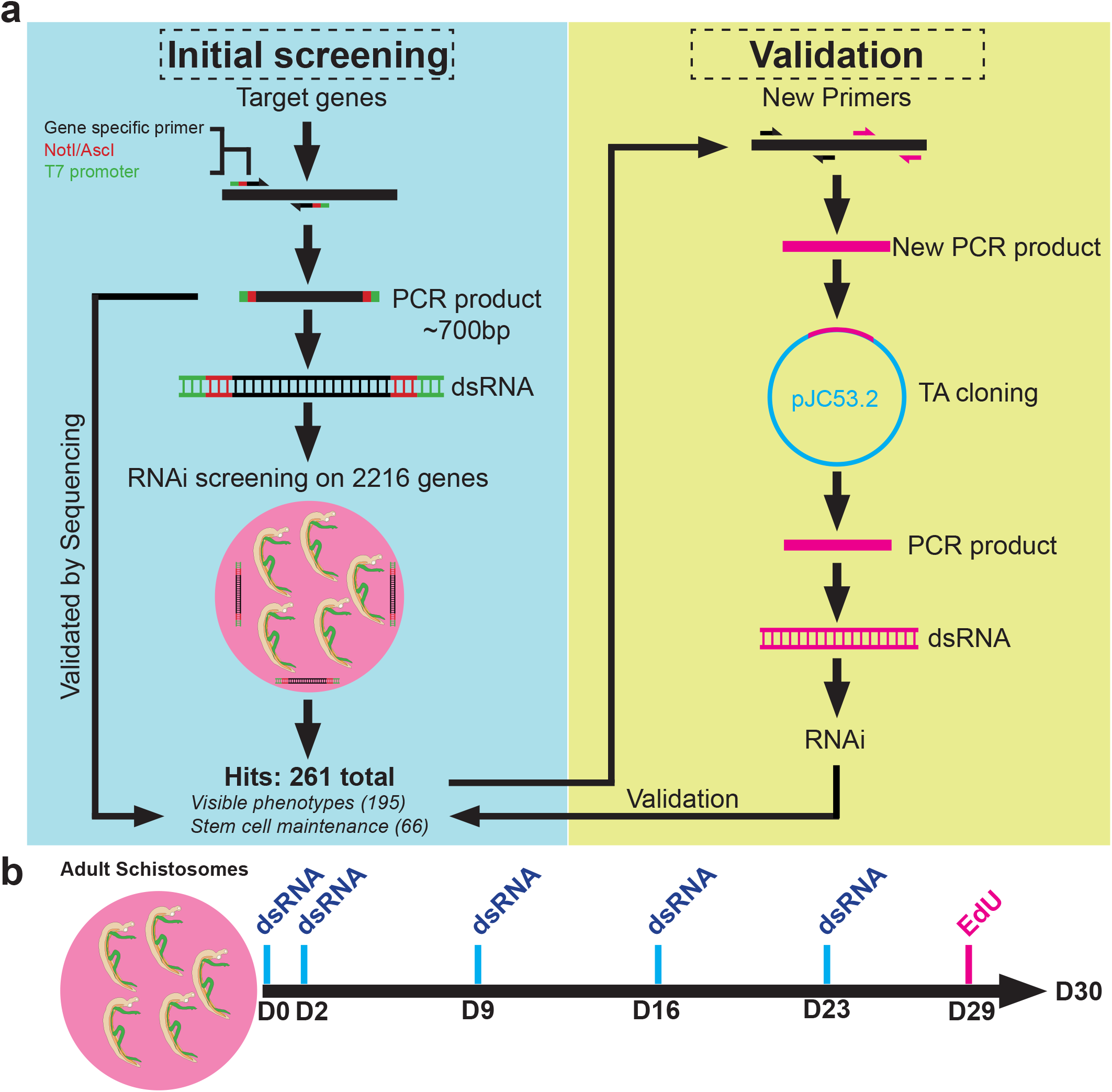
Platform for large-scale RNAi screening in *S. mansoni*. **a.** Pipeline for large-scale RNAi screening in *S. mansoni*. **b.** Double-stranded RNA treatment regime over the course of the 30-day treatment period of adult worms. During the entire experiment parasites are monitored for visible abnormalities and at D29 EdU is added to media to label proliferative cells in the parasites.

These parasites live in the veins surrounding the host intestines, and attachment to the vascular endothelium is essential *in vivo* for parasites to be kept from being swept away in the blood and trapped in host organs. Since detachment from an *in vitro* tissue culture substrate has been shown to precede more deleterious phenotypes^7^, and since under our *in vitro* culture conditions, healthy parasites firmly attached to the substrate using a combination of their oral and ventral suckers (**Supplementary Video 1**), we reasoned that substrate attachment would be a useful quantitative metric to define RNAi treatments that affect parasite vitality and predict *in vivo* survival. Therefore, during our 30-day experiments we monitored parasites every 48 hours for substrate attachment and any other visible defects. Schistosomes possess adult somatic stem cells, called neoblasts, that rejuvenate key parasite tissues, including the intestine and tegument (skin)^6,8^, and are likely to be essential for long-term parasite survival in the blood. The parasites also contain large numbers of proliferative germline stem cells (GSCs) in their reproductive organs^8^ which are essential for producing eggs that represent the central driver of parasite-induced pathology *in vivo^9^*. Therefore, we also monitored the maintenance of neoblasts and GSCs by labeling with the thymidine analog EdU prior to the conclusion of the experiment (**Fig. 1b**). Due to the variable rate at which the reproductive organs of female worms degenerate during *in vitro* culture^10^, stem cell proliferation was only monitored in male worms. At the conclusion of this initial screen we performed two major quality control steps for RNAi treatments resulting in attachment- or stem cell-related phenotypes. First, we confirmed the identity of every gene producing a phenotype by DNA sequencing. Second, where possible, we examined the specificity of our RNAi knockdown by designing new oligonucleotides targeting a non-overlapping region of genes that produced phenotypes (**Fig. 1a**, **Extended Data Fig. 1)**. To be considered a “hit” a gene must have shown a fully penetrant phenotype in three independent experiments. These studies identified 195 genes that were essential for parasite attachment, and thus potentially essential for worm survival *in vivo*. In addition to facilitating parasite substrate attachment, we also observed that 121 of these 195 genes were associated with other visible phenotypes including tissue and intestinal edema (36), head (26) and/or tegument (78) degeneration, muscular hypercontraction (6), and complete cessation of movement (death) (36) (**Fig. 2a**, **Supplementary Table 2**). In addition to these genes, we found that RNAi of an additional 66 genes resulted in stem cell maintenance defects but caused no other visible phenotypes (*e.g.*, substrate attachment) suggesting a selective role in stem cell maintenance (**Supplementary Table 3, Extended Data Fig. 2**).

**Fig2.**
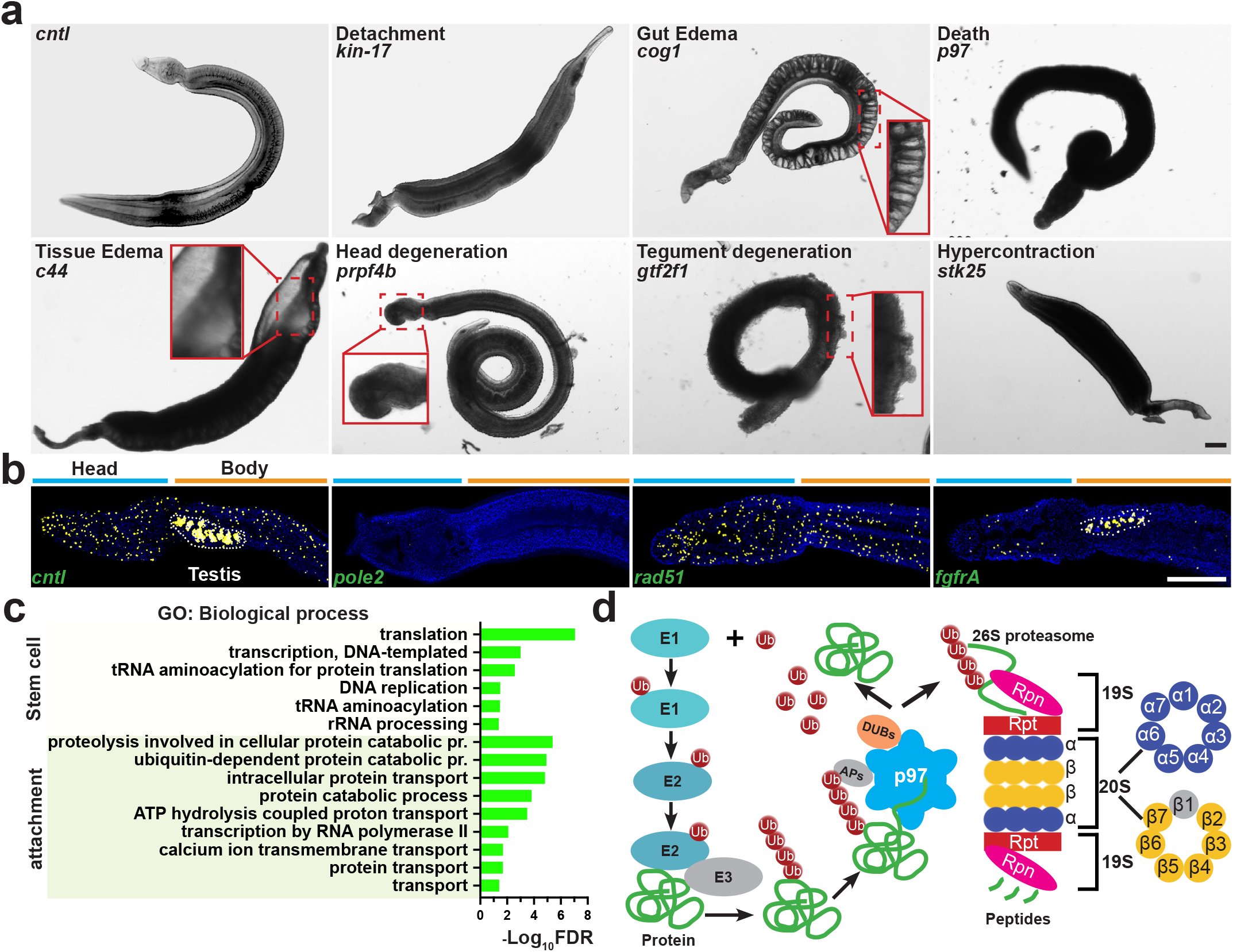
Summary of RNAi phenotypes. **a.** Categories of RNAi phenotypes observed. *kin-17* (Smp_023250), *cog* (Smp_132980), *p97* (Smp_018240), *c44* (Smp_136260), *prpf4b* (Smp_068960), *gtf2f1* (Smp_088460), *stk25* (Smp_096640). **b.** EdU-labeling (yellow) showing proliferative cells in somatic tissues and the testes. RNAi of DNA polymerase epsilon subunit (Smp_124120) leads to loss of all proliferative cells, whereas *rad51* (Smp_124230) or *fgfrA* (Smp_175590) lead to a selective reduction in the testes and soma, respectively. **c.** Gene Ontology analysis examining the biological processes of genes required for either stem cell maintenance or substrate attachment. **d.** A large fraction of genes resulting in visible phenotypes were associated with components of the Ubiquitin Proteasome System (UPS). Left, cartoon of the UPS. Colored UPS components correspond to genes associated with visible phenotypes. Scale bars: a, 100 μm; b, 200 μm

Of the 66 genes essential for stem cell survival over 90% (60/66) led to defects in the maintenance of both neoblasts and proliferative cells in the male testes (**Extended Data Fig. 2**). However, in a minority of cases some genes appeared to play more significant roles in maintaining proliferative cells in either the male germ line (*e.g.*, a RAD51 homolog) or the neoblasts (e.g., *fgfrA*, a previously-described FGF receptor homolog^8^) (**Fig. 2b**). In addition to genes necessary for cell cycle progression (e.g., *polo-like kinase)*, Gene Ontology enrichment analysis highlighted genes important for protein translation, including gene products involved in ribosomal structure, tRNA aminoacylation, and rRNA processing as important regulators of proliferative cell maintenance (**Fig. 2c**, **Extended Data Fig. 3**). Although this could reflect an enhanced sensitivity of actively proliferating cells to alterations in protein translation, recent work has highlighted “non-housekeeping” roles for translational regulators in controlling stem cell function^11^. Thus, it is worth exploring whether specific roles for translational control exist for regulating schistosome stem cell function.

Similar to previous whole organism large-scale RNAi studies in other metazoa^12,13^, we found that a large fraction of the 195 genes essential for parasite vitality (attachment) share sequence similarity (BLAST *E*-value < 1e-5) with genes in other organisms including *C. elegans* (91%), *Drosophila* (93%), the planarian *Schmidtea mediterranea* (97%), and humans (93%) (**Supplementary Table 4**). Some of these 195 schistosome genes with detachment phenotypes have *C. elegans*/*D. melanogaster* orthologs that lack any phenotypes (**Supplementary Table 5**); such genes could regulate novel schistosome-specific biology or represent opportunities for studies of *S. mansoni* to shed light on the function of poorly characterized animal gene families. Further examination of genes with attachment phenotypes by Gene Ontology analyses revealed that although this dataset was enriched for genes encoding regulators of protein transport and mRNA transcription (**Fig. 2c**, **Extended Data Fig. 3**), the dominant group of enriched genes were those encoding components necessary for protein turnover via the ubiquitin-proteasome system (UPS) (**Fig. 2c**, **Extended Data Fig. 3**). RNAi and pharmacological studies have implicated proteolysis by the proteasome as important for larval, and, more recently adult viability *in vitro^14,15^*. However, our data points to a much broader requirement for UPS components in these worms. Indeed, inspection of our RNAi dataset found that key components from virtually every arm of the UPS were required for adult parasite vitality during *in vitro* culture including: E1/E2 ubiquitin ligases and Deubiquitinating Enzymes (DUBs), the AAA-ATPase p97 that delivers proteins to the proteasome^16^, and nearly all regulatory and catalytic subunits of the proteasome complex^17^ (**Fig. 2d**). Indeed, RNAi of nearly all of UPS components resulted in extensive tissue degeneration and in some cases (e.g., *p97(RNAi)*) adult parasite death (**Extended Data Fig. 4**). Taken together, these data suggest that disruption not just of proteasome function, but any critical UPS components, results in reduced schistosome vitality *in vitro*.

To determine if any genes associated with attachment phenotypes encoded proteins targeted by existing pharmacological agents, we performed a combination of manual searches of the literature and bioinformatic comparisons with the ChEMBL database^18^ (**Supplementary Table 6**). This analysis uncovered 205 compounds potentially targeting 49 *S. mansoni* proteins. To gauge the utility of this approach to prioritize compounds with activity on adult parasites, we selected 14 of these compounds (**Supplementary Table 7**), including: FDA-approved drugs (*e.g.*, Ixazomib, Panobinostat), drugs currently or previously explored in clinical trials (*e.g.*, CB-5083, HSP990), or experimental compounds with activity in rodent models of disease (e.g., Thapsigargin, NMS-873). We then examined their activities on worms cultured *in vitro* using an automated worm movement tracking platform^19^ and by measuring the effects on parasite attachment following drug treatment. This analysis found that more than half of the compounds tested (8/14) on worms at 10 μM reduced parasite movement >75% and half of the compounds (7/14) caused fully penetrant substrate attachment defects by D7 post-treatment (**Fig. 3a-b**, **Supplementary Video 2**). Among the compounds that emerged from these studies was simvastatin, an HMG-CoA reductase inhibitor, that was previously shown to have effects on parasites both *in vitro* and *in vivo^20^*. We also evaluated these compounds on post-infective larvae (schistosomula), observing that 7 had profound effects on parasite movement (**Supplementary Table 8**), suggesting the potential of these compounds to target multiple schistosome life-cycle stages. Consistent with our observation that UPS function is critical for schistosome vitality (**Fig. 2d-f**), we found that the proteasome inhibitor ixazomib caused profound effects on both schistosome movement (**Fig. 3a**) and attachment (**Fig. 3b**), mirroring a recent report using the proteasome inhibitor bortezomib^14^. However, among compounds with the most potent effects on adult parasites were inhibitors of the UPS component p97: CB-5083, an ATP-competitive inhibitor^21^, and NMS-873, an allosteric inhibitor^22^, that both had sub-micromolar effects on adult worm movement (EC_50_ = 0.93 μM for NMS-873 and 0.16 μM for CB-5083) (**Extended Data Fig. 5**). Similar to the death observed following long-term *p97* RNAi treatment (**Fig 2a**), both NMS-873 and CB-5083 led to death *in vitro* (**Supplementary Video 3**). Despite their differing mechanisms of p97 inhibition (ATP-competitive vs allosteric), we noted similar deformations in the structure of the parasite tegument following treatment with either CB-5083 and NMS-873, suggesting that these compounds have similar pharmacological effects on the parasite (**Fig. 3c**). Given the prominent role for the UPS in schistosomes (**Fig. 2c-d**), we assessed if NMS-873 and CB-5083 affected UPS function by measuring the accumulation of ubiquitinated proteins using an antibody that recognizes K48 polyubiquitinated proteins marked for proteasome-mediated destruction^23^. Not only did we observe the accumulation of polyubiquitinated proteins following RNAi of *p97*, treatment of schistosomes with either CB-5083 or NMS-873 enhanced anti-K48 polyubiquitin labeling (**Fig. 3d**). We observed similar accumulation of polyubiquitinated proteins following either RNAi of *proteasome subunit beta type-2* or treatment with ixazomib (**Extended Data Fig. 5**). These effects on the degradation of ubiquitinated proteins appeared to be specific to inhibition of UPS function, rather than a non-specific effect due to reduced worm vitality, as treatment with the sarco/endoplasmic reticulum Ca^2+^-ATPase inhibitor thapsigargin, which also caused profound effects on worms (**Fig 3a, 3b**), did not alter the accumulation of polyubiquitinated proteins (**Extended Data Fig. 5**).

**Fig3.**
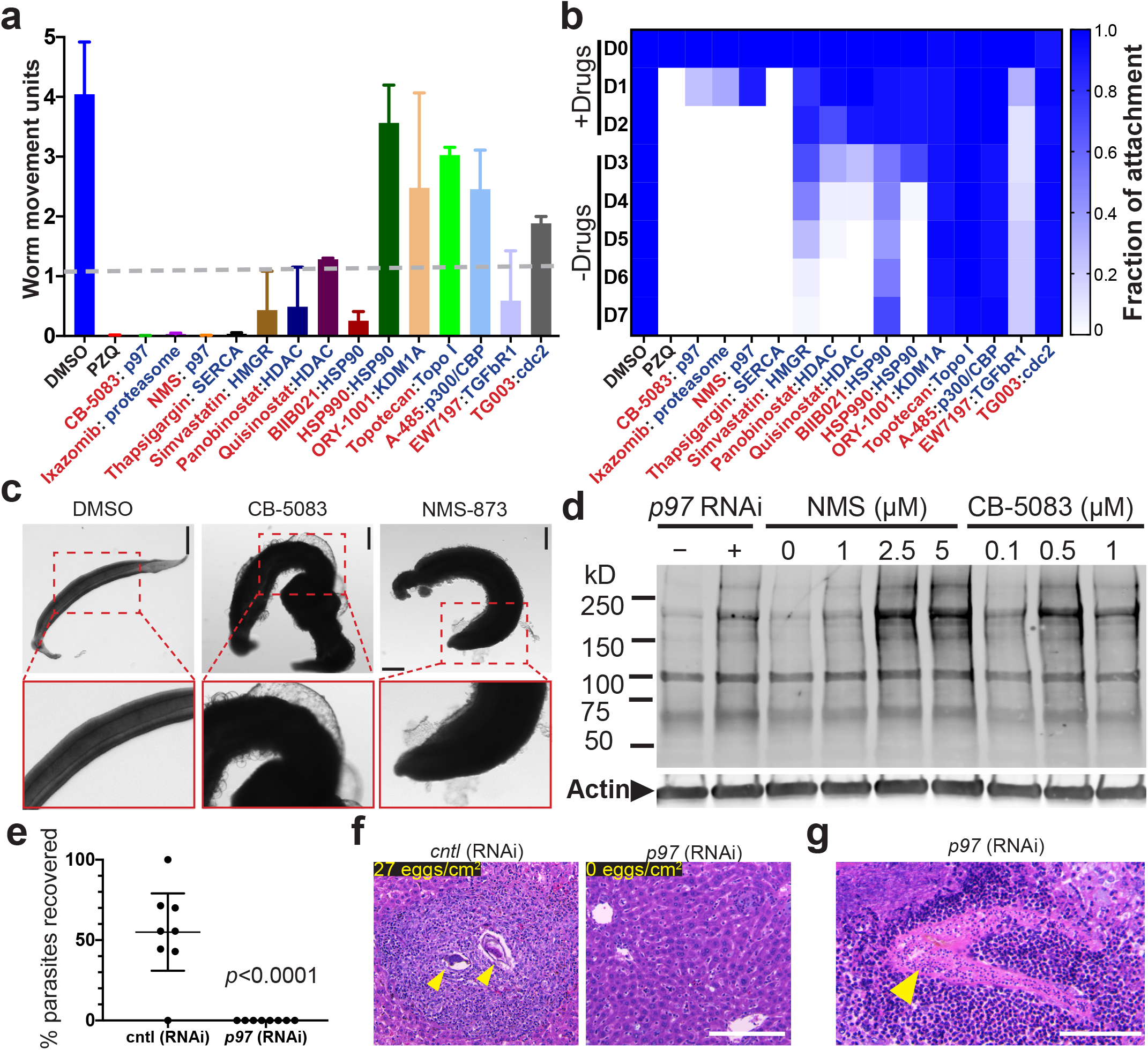
Compounds prioritized from RNAi studies have effects on schistosomes *in vitro*. **a.** Compounds (red text) predicted to target schistosome proteins (blue text) essential for parasite vitality from RNAi studies were examined at 10 μM for their effects on worm motility. Parasites were incubated with compounds and movement assessed after 72 hrs. Praziquantel (PZQ, 10 μM in 0.1% DMSO) and DMSO (0.1%) were used as positive control and negative controls, respectively. Dashed line shows threshold for 75% reduction in worm motility. Error bars represent standard deviation of the mean motility scores. *n* = 12 (three biological replicates, each compound was tested in duplicate - each replicate containing a pair of adult worms/well). **b.** Heatmap showing time course measuring the fraction of male worms attached to the substrate over a 7-day period following treatment of worms with compounds for 72 hours as in **a**. **c.** Treatment with either CB-5083 or NMS-873 at 10 μM (72 hrs) caused severe blebbing and delamination of the tegument. **d.** Western Blot for K-linked polyubiquitinated proteins. RNAi of *p97* or treatment of worms with p97 inhibitors caused an increase in the accumulation of polyubiquitinated proteins. Representative from 3 experiments. **e**, Percent recovery of male parasites treated with dsRNA specific to *p97* (Smp_018240; n = 8 transplants) or an irrelevant dsRNA (*control*; n = 8 transplants) following surgical transplantation of parasites into mice. ****, *p* < 0.0001, t-test **f.** Hematoxylin and Eosin staining of livers from recipient mice that received either control or *p97(RNAi)*. Schistosome egg-induced granulomas in livers were observed in control RNAi recipient mice, but not in *p97(RNAi)* recipient mice. Counts of eggs per liver section are shown in top left, *n=3*. **g.** Transplanted parasites from *p97(RNAi)* treatments were found trapped and in various stages of degeneration in livers of recipient mice. Scale bar: c, f, g, 100 μm.

To determine if UPS function is broadly required for adult schistosomes *in vivo*, we depleted UPS components using RNAi and surgically transplanted these worms into the mesenteric veins of recipient mice (**Extended Data Fig. 6**) to measure parasite egg deposition in host tissues and parasite survival^7^. Following hepatic portal perfusion, we recovered about 55% of control RNAi-treated worms originally transplanted (**Fig. 2e**, **Extended Data Fig. 6**) and these parasites established patent infections depositing large number of eggs into the livers of recipient mice (**Fig. 2f**, **Extended Data Fig. 6**). In contrast, we failed to recover parasites following hepatic portal perfusion from mice receiving *p97* (**Fig. 2e**) or *proteasome subunit beta type-2* (**Extended Data Fig. 6)** RNAi-treated worms. Additionally, the livers in these mice were devoid of eggs, as a consequence, we observed no signs of egg-induced granulomas (**Fig. 2f**, **Extended Data Fig. 6**). We did, however, observe RNAi-treated parasites at various stages of being infiltrated by host immune cells in the livers of recipient mice (**Fig. 2g**, **Extended Data Fig. 6**), suggesting these parasites are unable to remain in the portal vasculature and are cleared via the immune system in the liver. Thus, several components of the UPS are essential for schistosome survival *in vivo*. Recent studies from a variety of human parasites have highlighted the potential for therapeutically targeting UPS function by inhibition of the proteasome^14,24,25^. Our data suggest that targeting another critical (and druggable^21,22^) mediator of UPS function (*i.e.*, p97) may have therapeutic potential, not just against schistosomes, but against a variety of important human parasites.

Another prominent group of potentially druggable targets to emerge from our RNAi screen were protein kinases, 19 of which led to defects in either parasite attachment or stem cell maintenance. The most striking protein kinase-related phenotypes resulted from RNAi of two STE20 serine-threonine kinases: *tao* and *stk25*, which are homologs of the human TAO1/2/3 and STK25/YSK1 protein kinases, respectively. RNAi of either of these kinases led to rapid detachment from the substrate (**Extended Data Fig. 7**) and a concomitant posterior paralysis and hypercontraction of the body, such that the parasites were shorter than controls and took on a distinctive banana-shaped morphology (**Fig 4a-b**, **Supplementary Video 4**). Aside from RNAi of *stk25* and *tao*, this banana-shaped phenotype was unique, only observed in our screening following RNAi of a CCM3/PDCD10 homolog (Smp_031950), a known heterodimerization partner with the mammalian STK25 kinase^26^. We failed to observe death of either *stk25-* or *tao-* depleted parasites during *in vitro* culture; however, following surgical transplantation we noted a significant reduction in the recovery of *tao* or *stk25* RNAi-treated parasites from recipient mice and these recipient mice displayed little signs of egg-induced granuloma formation (**Extended Data Fig. 7**). Thus, both *tao* and *stk25* appear to be essential for schistosome survival *in vivo*.

**Fig4.**
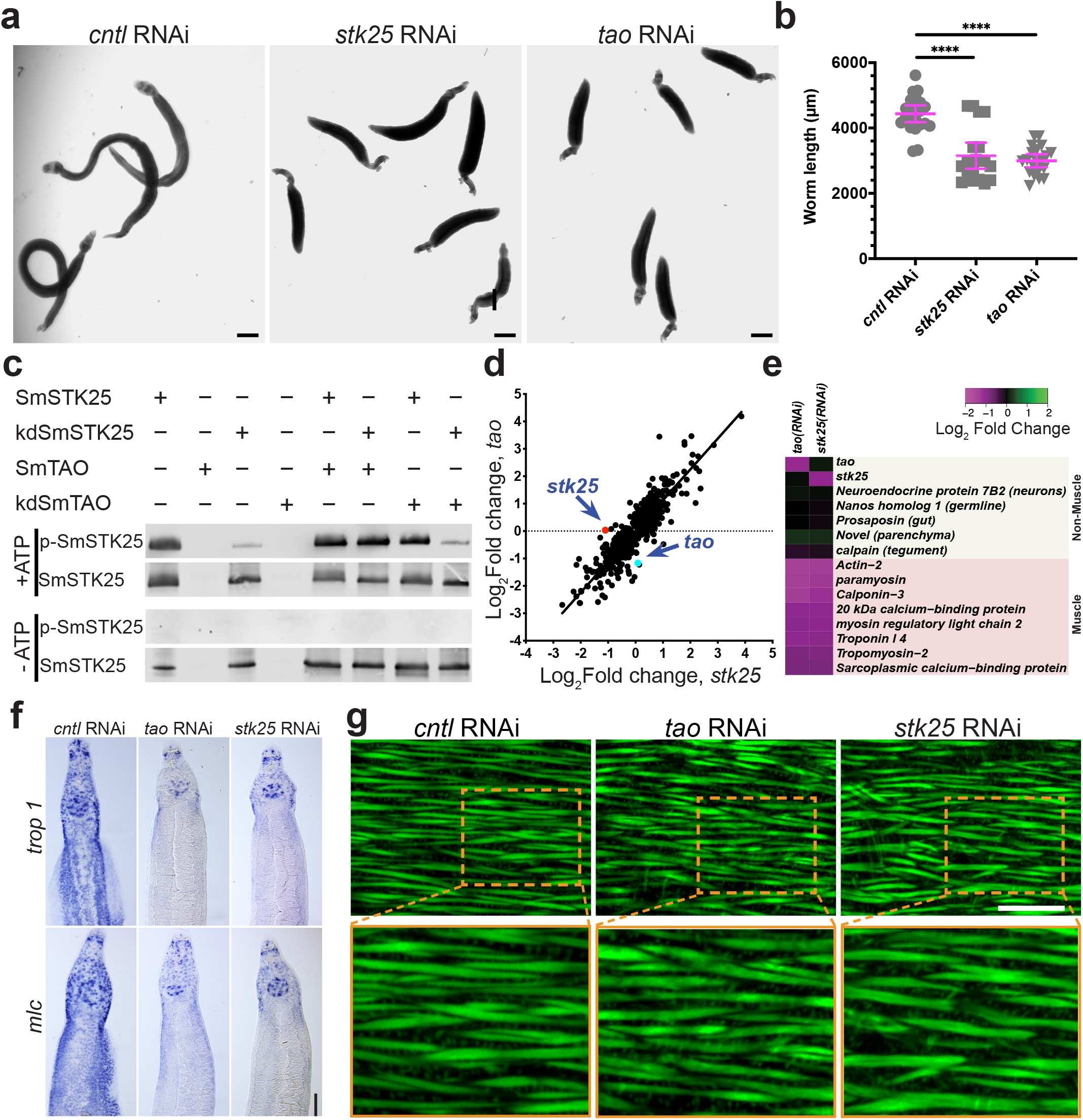
The protein kinases SmSTK25 and SmTAO are essential to maintain muscular function. **a.** RNAi of *stk25* or *tao* causes parasites to become hypercontracted. **b.** *stk25* and *tao* RNAi-treated parasites are shorter than control RNAi-treated worms. >19 parasites monitored over 4 experiments. *p* < 0.0001, t-test. **c.** Western blot to detect phosphor-T173 (p-SmSTK25) or total SmSTK25 following an *in vitro* kinase reaction with recombinant proteins in the presence or absence of ATP. Active SmSTK25 can autophosphorylate T173, as can SmTAO when incubated with kinase dead SmSTK25 (kdSmSTK25). T173 phosphorylation was dependent on ATP. kdSmTAO represents kinase dead SmTAO. Representative of 2 experiments. **d.** Dot plot showing the relationship between the differentially expressed genes following either *stk25* or *tao* RNAi-treatment. These expression profiles were highly correlated (*R* = 0.9, *p* < 0.0001). **e.** Heatmap showing that many muscle-specific transcripts were down-regulated following RNAi of *tao* or *stk25*. **f.** *in situ* hybridization to detect the expression of *tropomyosin 1* (Smp_340760) and a myosin light chain (Smp_132670) following RNAi of *tao* or *stk25* at D13. **g.** Phalloidin labeling to mark F-actin in muscle cells of RNAi treated parasites at D13 indicating that muscle fibers are intact at this timepoint after depletion of *tao* or *stk25*. Scale bars: a, 500 μm; f, 100 μm; g, 20 μm.

Given the unique and specific nature of the *stk25* and *tao* associated “banana” phenotype we reasoned that these kinases may be acting in concert to mediate similar signaling processes in the worm. Recent data suggests that the *Drosophila* STK25 ortholog (GCK3) is a substrate of TAO and that these proteins function in a signaling cascade essential for tracheal development^27^. Consistent with these studies, we too observed that recombinant *S. mansoni* STK25 (SmSTK25) could serve as a substrate for the *S. mansoni* TAO (SmTAO) in an *in vitro* kinase assay (**Extended Data 8**). The human STK25 is activated by phosphorylation of a conserved threonine residue within its activation loop^28^. By mass spectrometry we observed that this conserved threonine within the predicted SmSTK25 activation loop (T^173^) was phosphorylated following incubation of recombinant SmTAO with catalytically inactive SmSTK25 in the presence of ATP (**Extended Data 8**). To explore this observation in more detail we performed western blotting on *in vitro* kinase reactions using an antibody that recognizes phosphorylation of the conserved threonine in the activation loop of vertebrate and invertebrate STK25 orthologs^27^. Validating the specificity of this antibody of against phosphorylated T^173^ on SmSTK25, we detected robust SmSTK25 T^173^ autophosphorylation following an *in vitro* kinase reaction; this signal was abrogated when ATP was omitted from the reaction or when the SmSTK25 catalytic K^48^ residue was mutated to R (**Fig. 4c**, **Extended Data 10**). Consistent with our mass spectrometry results, we detected robust phosphorylation of T^173^ when recombinant SmTAO was incubated with kinase dead SmSTK25 (**Fig. 4c**), suggesting that SmTAO can phosphorylate a residue key for the activation of the mammalian STK25.

Given their phenotypic similarities and our biochemical observations, we reasoned that the schistosome TAO and STK25 might be acting in a signaling module to mediate critical processes in the parasite. To define these processes, we performed transcriptional profiling on RNAi-treated parasites just prior to the timepoint in which we observed detachment and hypercontraction (Day 6 and Day 9 for *tao* and *stk25* RNAi treatments, respectively) (**Extended Data Fig. 9**). We reasoned that transcriptional changes common to both *stk25* and *tao* RNAi data sets would provide details about any processes regulated by these proteins. Consistent with the model that these kinases cooperate in the parasite, we found that expression of differentially regulated genes following RNAi of either *tao* or *stk25* were highly correlated (**Fig. 4d**) and that more than half of these differentially regulated genes were common in both datasets (**Extended Data Fig. 9, Supplementary Table 9**). Importantly, RNAi of either *tao* or *stk25* was specific, not affecting expression of the other kinase gene of this pair (**Fig. 4c, d**). To better understand the genesis of the phenotype associated with loss of *tao* or *stk25*, we examined the tissue-specific expression of differentially-regulated genes on an adult schistosome single cell expression atlas using cells from schistosome somatic tissues^29^. Strikingly, we found that roughly 40% (51/129) of the most down-regulated genes following *tao* and *stk25* RNAi (Log_2_ Fold Change < −0.5, adjusted *p*-value < 0.00001) were specific markers of parasite muscle cells (**Extended Data Fig. 10, Supplementary Table 9**). Indeed, nearly half of all mRNAs specifically-enriched in muscle cells (60/135) from this single cell atlas, including key muscle contractile proteins (*e.g*, Troponin subunits Actins, Myosin light/heavy chains, and Tropomyosin), were significantly down-regulated following RNAi of both *tao* and *stk25* (**Fig. 4e**, **Extended Data Fig. 10, Supplementary Table 10**). Importantly, these transcriptional effects appeared to be largely specific to parasite muscles, since comparatively few markers specific to other major somatic organ systems (neurons, gut, parenchyma) were affected by RNAi of these kinases (**Fig. 4e**, **Extended Data Fig. 10, Supplementary Table 10**). In principle, loss of muscle-specific mRNAs could be due to either loss of muscle cells or down-regulation of muscle-specific mRNAs. To distinguish between these possibilities, we performed labeling with phalloidin to mark F-actin in schistosome muscle fibers and *in situ* hybridization to detect muscle-specific mRNAs. Within a few days of RNAi-treated parasites adopting their banana-shaped phenotype, we noted a dramatic reduction in the expression of mRNAs encoding the contractile proteins Tropomyosin 1 and a Myosin Light Chain by *in situ* hybridization (**Fig. 4f**), but observed no major qualitative defects in phalloidin labeling in the muscle fibers within anterior or posterior body wall muscles (**Fig 4g**, **Extended Data Fig. 8**). Thus, it appears that these kinases are required to maintain the transcription of a large number of muscle-specific mRNAs in intact muscle cells. Interestingly, we noted that the heads of *tao* and *stk25* RNAi parasites, which retained their capacity for movement (**Supplementary Video 4**), partially maintained the expression of muscle-specific mRNAs (**Fig. 4f**). Thus, there appears to be a relationship between the maintenance of muscle-specific mRNA expression and locomotion. Taken in their entirety, our data are consistent with the model that STK25 and TAO kinases cooperate (perhaps with TAO directly activating STK25) in the schistosome to mediate a signaling cascade essential for sustaining transcription of muscle-specific mRNAs. As a consequence, loss of either SmSTK25 or SmTAO activity results in muscular function defects and this compromises parasite survival *in vivo*. Although the essentiality of the three mammalian TAO homologs is unclear, whole body knockouts of mouse STK25 are homozygous viable displaying no obvious deleterious phenotypes^30^. Thus, SmSTK25 function appears to be a schistosome-specific liability for survival when compared to mammals. Given this, and the druggable nature of kinases, we suggest that SmSTK25 represents a high-value target for therapeutic intervention.

Technological advances have paved the way for large-scale analyses of gene function in protozoan parasites^31–33^, but, unfortunately, comparable resources have not yet materialized for any helminth parasite. Here, we have performed the largest systematic analysis to date of gene function in schistosomes, examining roughly 20 percent of the protein coding genes in the parasite. Our RNAi studies, together with bioinformatics, have allowed us to effectively prioritize targets essential in vivo (e.g., STK25, TAO, and p97) and potential specific inhibitors with *in vitro* activities on worms (**Fig 3a-b**). Thus, future efforts should not only explore compounds our bioinformatic approaches have already uncovered (**Supplementary Table 6)**, but also larger libraries of compounds with known molecular targets (e.g., the REFRAME collection^34^). Such studies are likely to be an efficient means to identify existing drugs for potential repurposing against schistosomes. Not only does this study enhance our understanding of schistosome biology, and serve as a template for conducting further genome-scale studies of schistosome gene function, it provides a new lens to prioritize genes of interest in other medically- and agriculturally-important parasitic flatworms (e.g., tapeworms and flukes). Collectively, we anticipate this study will expedite the discovery of new anthelmintics.

## Supplementary Information Guide

**Supplementary Table 1.** Information of 2,320 genes selected for RNAi screening.

**Supplementary Table 2.** Details of 195 genes with detachment phenotypes.

**Supplementary Table 3.** Details of 66 genes with phenotypes in EdU incorporation.

**Supplementary Table 4.** Similarity of schistosome genes with phenotypes with gene products from other organisms by BLAST.

**Supplementary Table 5.** *S. mansoni* genes with detachment phenotypes, whose *C. elegans* and *D. melanogaster* orthologs lack phenotypes in WormBase/FlyBase

**Supplementary Table 6.** Human homologs of *S. mansoni* RNAi hits and their potential inhibitors.

**Supplementary Table 7.** Details of 14 selected inhibitors to test on schistosome.

**Supplementary Table 8.** Evaluation of compound activity on schistosomula

**Supplementary Table 9.** Analysis of transcriptional changes following *stk25* or *tao* RNAi treatment by DESeq. Second tab shows which somatic cell clusters the most down-regulated (*p*<0.00001 Log_2_ Fold Change <−0.5) genes following *stk25* or *tao* RNAi treatment are expressed.

**Supplementary Table 10.** Analysis of expression of somatic tissue-specific markers following *stk25* or *tao* RNAi treatment. Tissue-specific markers down-regulated (Log_2_ Fold Change < 0 and pAdj < 0.000001) following both *stk25* and *tao* RNAi-treatments are highlighted in red.

**Supplementary Video 1. Adult worms after 30 days of i*n vitro* treatment with control dsRNA.** On day 30, worms were physically active and firmly attached to the bottom of the dish.

**Supplementary Video 2. Parasites died in 72 hours *in vitro* after inhibition by CB-5083 or NMS-873**. Adult worms were dead following 72 hours of treatment with 1 μM CB-5083 or 5 μM NMS-873. Tegmental damage was observed on these worms. DMSO was used as a control.

**Supplementary Video 3. *stk25/tao* RNAi treated worms become hypercontracted and paralyzed by D13 following dsRNA treatment.**

**Supplementary Video 4. Effects of various compounds on parasites.** Parasites were treated with compounds for 72 hours at a concentration of 10 μM. We observed various worm defects ranging from death, to tissue degeneration, and detachment from the substrate. DMSO and PZQ were used as negative and positive controls, respectively.

## Acknowledgements

We thank Megan McConathy, Caroline Furrh, and Dr Giampaolo Pagliuca for technical assistance and Fiona Hunter and Nicolas Bosc for advice about retrieving data from ChEMBL. Mice and *B. glabrata* snails were provided by the National Institute of Allergy and Infectious Diseases (NIAID) Schistosomiasis Resource Center of the Biomedical Research Institute (Rockville, MD, USA) through National Institutes of Health (NIH)-NIAID Contract HHSN272201000005I for distribution through BEI Resources. The work was supported by the National Institutes of Health R01AI121037 (JJC), the Welch Foundation I-1948-20180324 (JJC), the Burroughs Wellcome Fund (JJC), and the Wellcome Trust 107475/Z/15/Z (JJC/KFH/MB).

## Methods

### Parasites

Adult *S. mansoni* (NMRI strain) (6–7 weeks post-infection) or juvenile worms (4–5 weeks post-infection) were harvested from infected mice by hepatic portal vein perfusion with 37°C DMEM (Mediatech, Manassas, VA) plus 8% Horse Serum and heparin. Parasites were rinsed in DMEM + plus 8% Horse Serum and cultured (37°C/5% CO_2_) in Basch’s Medium 169^35^ and 1× Antibiotic-Antimycotic (Gibco/Life Technologies, Carlsbad, CA 92008). Experiments with and care of vertebrate animals were performed in accordance with protocols approved by the Institutional Animal Care and Use Committee (IACUC) of UT Southwestern Medical Center (approval APN: 2017-102092).

### Initial RNAi screening

Primers were designed to amplify ~700 bp fragment using BatchPrimer3 http://batchprimer3.bioinformatics.ucdavis.edu/index.html. For genes shorter than 700 bp, primers were designed to cover as much of the transcript as possible. For reverse transcription of double stranded RNAs, a T7 promoter sequence (GAATTTAATACGACTCACTATA) was added to the 5’ end of each oligo. To facilitate DNA sequencing of cDNAs associated with RNAi phenotypes, we added a *NotI* or *AscI* restriction enzyme site between the T7 and gene-specific sequences on the forward and reverse oligos, respectively. These oligos were synthesized and packaged in 96-well plates and used for PCR using adult schistosome cDNA as a template. 5 μL of PCR products were then used for *in vitro* transcription to generate dsRNA in 100 μL as previously described^36^. After overnight incubation at 37 °C, dsRNAs were annealed by a successive 3-min gradient incubation at 95 °C, 75 °C, and 55 °C, then cooled down at room temperature for 5 min. The presence and size of PCR products and dsRNA were all analyzed by agarose gel electrophoresis and samples stored at −20 °C. For RNAi treatments, approximately 5 pairs of adult parasites were placed in 3 mL Basch 169 media in a 12-well plate and treated with 20 μL dsRNA at D0, D2, D9, D16 and D23. To examine cell proliferation, the media were supplemented with EdU (10 μM) at D29. On day 30, videos were captured for RNAi treatments that caused visible RNAi phenotypes and after video acquisition all parasites were fixed and processed for EdU detection^8^. During the entire 30D RNAi treatment regime, media was changed every 1-2 days and worm attachment and morphological changes were monitored. Videos RNAi treatments causing visible phenotypes can be found at: https://datadryad.org/stash/share/R4pxckHwhrBqUyfMkuH2FHhRJzdjP_wKLbkZpVCP8QE.

To validate hits from the initial RNAi screening, the original PCR products were digested with NotI (NEB) for 30 min at 37 °C, gel purified (Zymoclean Gel DNA Recovery Kit), and sequenced with a T7 primer. Sequences of genes validated by sequencing were uploaded into BatchPrimer3 to design new primers that amplify a fragment sharing no overlap with the PCR products from the initial RNAi screening. In cases where genes’ sequences were too short to design new oligos, we retained the original primer sequences. These primers were synthesized without further modification, used to generate PCR products, and then inserted into pJC53.2 using TA cloning^36^. These plasmids were purified from *E. coli*, validated by sequencing, and used as a template to generate dsRNA. We then repeated the RNAi treatment regime used in the original screen.

### Parasite labelling and imaging

Whole-mount in situ hybridization^6^, EdU detection^8^, and phalloidin staining^37^ were performed as previously described. For in situ hybridization, riboprobes were generated from cDNA fragments amplified using primers for tropomyosin-1 (Smp_340760, gagaaagagaatgctatggaaagagc/cctcattttgtagtttagatacttgacg) or myosin light chain (Smp_132670, gttgctctgtgttaagttaacatggg/gttagtcctaaatgtcttgattgcc). Brightfield images of in situ hybridizations and worm morphology/movement were imaged using a Zeiss AxioZoom V16 (Zeiss, Germany) equipped with a transmitted light base and a Zeiss AxioCam 105 Color camera. Fluorescent images were acquired using a Nikon A1+ laser scanning confocal microscope.

### Transplantation of dsRNA-treated Schistosomes

Parasites 4–5 weeks post-infection were recovered from mice and treated with 30 μg/ml dsRNA for 4 days in Basch Media 169 with a daily replacement of media and dsRNA and surgically transplanted into naïve mice as previously described^7^. On day 26 post-transplantation, mice were sacrificed and perfused to recover parasites. Male and female parasites were counted and livers were removed and fixed for 30–40 hours in 4% formaldehyde in PBS. The percentage of parasite recovery was determined by dividing the number of male worms transplanted by the number of male parasites recovered following perfusion. Livers from individual mice were sectioned and processed for Hematoxylin and Eosin staining by the UT Southwestern Molecular Pathology Core.

### Detection of polyubiquitinated proteins by western blot

For RNA interference, 10 single-sex male adult worms (6 weeks post infection) were treated with 30 μg/mL dsRNA in Basch Media 169 for 8 days. Media and dsRNA were changed daily. On day 9, worms were collected and flash frozen. For drug treatment, 10 male adult worms (single or paired with females) were supplemented with either DMSO, NMS 873 or CB 5083. After 24hrs, male parasites were separated from females using 0.25% tricaine in Basch Media 169 and flash frozen. Male worm samples were homogenized with a pestle in 50 μL lysis buffer containing 2 x sample buffer, protease inhibitor cocktail (Roche, cOmplete Mini, EDTA-free Tablets) and 10mM DTT. The lysates were then sonicated on high for 5 min (30 sec on, 30 sec off) using a Bioruptor UCD-200. Lysates were centrifuged for 5 min at 10,000 *g* to remove debris. Total protein was measured using the Detergent Compatible Bradford Assay (Pierce). 35 μg of protein samples denatured in SDS Sample buffer (95°C for 5min) were separated on a Bio-Rad 4-20% TGX Stain-Free gel along with Precision Plus Protein Dual Color Standards (Bio-Rad) as a marker. Proteins were then transferred to a nitrocellulose membrane (Bio-Rad) and confirmed by Ponceau S staining. The membrane was blocked in a 1:5 solution of Li-Cor Odyssey Blocking buffer in PBS for 1hr before being immunoblotted overnight at 4°C with 1:500 K48-linkage Specific Polyubiquitin Antibody (Cell Signaling Technology, 4289S) and 0.01 μg/mL mouse anti-actin antibody (Developmental Studies Hybridoma Bank, JLA20) diluted in a 1:5 solution of Li-Cor Odyssey Blocking buffer in PBS. The membrane was washed 3x in TBST and then incubated in 1:5 Li-Cor Odyssey Blocking buffer containing the secondary antibodies (1:10,000 Li-Cor, 925-68071, goat anti-rabbit IRDye 680 RD, and 1:20,000 Li-Cor, 925-32280, goat anti-mouse IgM IRDye 800CW) for 1hr at RT. The blot was washed in TBST 3x before being imaged on a Li-Cor Odyssey Infrared Imager.

### Compound prioritization

To manually search for existing drugs targeting “detachment” hits from our RNAi screen, we performed protein-protein BLAST against the *Homo sapiens* proteome to find the closest human homolog to our RNAi hits. We then manually searched a variety of databases (*e.g.*, genecards, google, DrugBank, Therapeutic Targets Database) and chemical vendors (*e.g.*, seleckchem) for inhibitors against these human proteins. In each instance, we consulted the published literature to give preference to compounds likely to be selective for a given target. If several such drugs were available, preference was given to those that were also orally bioavailable and/or FDA approved/in clinical trials. For larger-scale discovery of compounds, the *S. mansoni* protein sequences of genes with ‘detachment’ phenotypes were used to search the ChEMBL database^18^, to identify compounds predicted to interact with them. To do this, we followed the protocol previously described^38^ with the following differences. First, for each *S. mansoni* gene, we identified its top BLASTP hit among all ChEMBL targets, as well as any ChEMBL targets having BLAST hits with *E*-values within 10^5^ of the top hit’s *E*-value; and then extracted from ChEMBL the drugs/compounds with bioactivities against those particular ChEMBL targets. Second, when calculating the ‘toxicology target interaction’ component of a compound’s score, we checked whether ChEMBL predicted with probability >0.5 that the compound interacts with one of 108 toxicology targets curated from^39–41^.

### Evaluation of effects of compounds on worms

*in vitro* evaluation of selected compounds (single-point concentrations, 10 μM in 0.1% DMSO) on adult movement was replicated three times using a single worm pair per well (two technical replicates each time, n = 12), as previously described^42^. Worm pairs co-cultivated with DMSO (0.1% negative control) and Praziquantel (PZQ) (10 μM in 0.1% DMSO; positive control) were included in each experiment. Following incubation at 37°C for 72 hrs in a humidified atmosphere containing 5% CO_2_, a digital image processing-based system was used for the assessment of parasite motility. Both hardware and software components of this system (WormassayGP2) were inspired by the digital macroscopic imaging apparatus previously described^19^, with minor modifications to the source code (supporting USB video class, UVC, camcorders and the High Sierra MacOS) and user interface (allowing manual manipulations to recording duration). A dose-response titration (10 μM – 0.156 μM) of CB-5083 and NMS-873 was performed to assess adult worm anti-schistosomal potencies. Each titration point was performed in triplicate (a pair of worms for each replicate). Worm movement was recorded with WormassayGP2, as mentioned above. Mean motility scores were calculated for each titration point and dose-response curves were derived in comparison to worms co-cultured in DMSO (0.1% v/v; negative control; 100% motility) and PZQ (10 μM in 0.1% DMSO; positive control; 0% motility). Anti-schistosomula activities of the selected compounds were assessed using the high-content imaging platform Roboworm as previously described^42,43^. Compounds (reconstituted in dimethyl sulfoxide, DMSO; 10 mM stock concentration) were initially tested at two different concentration points (10 μM and 50 μM, in 0.625% DMSO) along with negative (0.625% DMSO) and positive controls (PZQ at 10 μM final concentration in 0.625 % DMSO). Schistosomula/compound co-cultures were then incubated at 37°C for 72 h in a humidified atmosphere containing 5% CO_2_ before phenotype and motility metrics were assessed. Two-fold titrations (10 μM, 5 μM, 2.5 μM, 1.25 μM and 0.625 μM) were subsequently conducted for all compounds consistently identified as hits at 10 μM in the primary screens. Single point schistosomula screens (10 μM) were repeated three times whereas dose response titrations were performed twice (in each screen two technical replicates were included). Phenotype and motility scores deriving from the titration of each compound were collected to generate approximate EC_50s_ using GraphPad Prism. To quantify adult worm attachment to the substrate following drug treatment, freshly perfused adult worms were sorted into a 6-well plate with 3 mL Basch 169 media and cultured overnight. The following day (D0) unattached worms removed and compounds were added to the media to a final concentration of 10 μM. Media and drug were replaced on D1 and D2. Media with no drug was added on D3 and D5. Parasite attachment was monitored from D0 to D7.

### RNAseq for *stk25* and *tao* RNAi-treated worms

To examine gene expression changes following loss of *tao* or *stk25*, 10 adult worm pairs were placed into 6-well plates and cultured in 3 mL Basch 169 supplemented with 30 μg/mL dsRNA for 3 days. Media and dsRNA were replaced daily. On D3, dsRNA-containing media was removed and worms were maintained in 6 mL Basch 169 media that was replaced every other day. On day 6 (*tao(RNAi)*) or D9 (*stk25(RNAi))* worms were anesthetized with 0.25% tricaine and separated by sex. As controls, worms cultured in parallel were treated with an irrelevant dsRNA^8^. For RNA extraction, male worms were collected, excess media removed, and 100 μL of TRIZOL was added. Parasites were then flash frozen in liquid N_2_, homogenized with a micro pestle, the volume of TRIZOL was brought to 600 μL before RNA was purified using a Zymo Direct-zol RNA miniprep kit and processed for Illumina sequencing. RNAseq data was mapped to the *S. mansoni* genome (v7) using STAR and differential expression was analyzed by DESeq2 as previously described^44^. To define correlations between genes differentially regulated following RNAi of *tao* and/or *stk25*, we compiled a list of all genes that had significantly changed expression in either *stk(RNAi)* or *tao(RNAi)* datasets and plotted their log2 fold-change expression in GraphPad Prism to calculate a Pearson’s correlation coefficient. To evaluate the effects of *tao* and *stk25* RNAi on gene expression in specific tissues and cell types we collapsed related cell types from a *S. mansoni* single cell atlas^29^ into 10 broad clusters of male somatic cell types (muscles, neurons, neoblasts, gut, etc.) (**Extended Data Fig. 9**). Genes highly enriched in these clusters were determined using Seurat v3.1.1^45^ (parameters = logfc.threshold = 1, min.pct = 0.5) and compared to genes down-regulated (*p adj* < 0.000001) following both *tao* and *stk25* RNAi.

### Purification of Recombinant STK25 and TAO

Baculovirus expressing wildtype *Schistosoma mansoni* Smp_068060 (TAO) or Smp_096640 (STK25) with a C-terminal His_6_ tag was generated by GenScript (Piscaaway, NJ). cDNA encoding kinase-dead versions of both kinases were subcloned into the pFastBac1 vector with C-terminal His_6_ tag and baculovirus was generated according to the manufacturer’s instructions using the Bac-to-Bac Baculovirus Expression System (Invitrogen). Baculovirus was used as a 3^rd^ pass virus to infect Sf9 cells grown in Gibco Sf 900 III SFM (Thermofisher Scientific) supplemented with 1% FBS and Antibiotic-Antimycotic solution (Sigma-Aldrich). Cells were harvested 72hrs post infection for SmSTK25 expression and 48hrs post infection for SmTAO expression. Frozen cell pellets were lysed with 20mM Tris, pH 8.0, 5mM MgCl_2_, 300mM NaCl, 1% Triton X-100 (Fisher Scientific), DNase (24μg/ml), 10% glycerol, 3mM 2-mercaptoethanol, and protease inhibitors (1μg/ml aprotinin, 2μg/ml leupeptin, 1mM benzamidine, and 0.2mM PMSF). After homogenization, the suspension was centrifuged for 1h at 186,000 x *g* and the supernatant was rotated with Ni^2+^-NTA resin (Qiagen) for 1.5hrs. The resin was washed, and the protein was eluted in 20mM Tris, pH 8.0, 5mM MgCl_2_, 300mM NaCl, 0.05% Triton X-100, 10% glycerol, 3mM 2-mercaptoethanol, 150mM Imidazole, pH 8.0 and protease inhibitors (1μg/ml aprotinin, 2μg/ml leupeptin, 1mM benzamidine, and 0.2mM PMSF). Eluted proteins were either flash frozen or further dialyzed overnight into storage buffer (20mM Tris, pH 8.5, 5mM MgCl_2_, 150mM NaCl, 0.5mM DTT, 10% glycerol, and 1mM benzamidine) and flash frozen to −80°C.

To generate an anti-SmSTK25 antibody, a C-terminal fragment of SmSTK25 corresponding to AA513-622 was amplified and sub-cloned into pET28 vector with a C-terminal His_6_ tag for expression in *Escherichia coli*. This fragment was purified from transformed Rossetta 2 cells grown in LB medium and induced with 1mM isopropyl 1-thio-β-D-galactopyranoside for 16 hrs at 18°C. Cells were pelleted and resuspended into lysis buffer containing 50mM Tris, pH8.0, 300mM NaCl, 10% glycerol, and protease inhibitors (0.2mM PMSF, 2μg/ml aprotinin, and 2μg/ml leupeptin). The suspension was freeze-thawed and the following reagents were added to a final concentration of 1mg/ml lysozyme, 1% Triton X-100, 5μg/ml DNase. After homogenization and sonication, lysate was centrifuged for 40 min at 186,000 x *g*, and rotated with Ni^2+^-NTA resin for 1.5 hrs. The resin was washed, and protein was eluted in lysis buffer containing 300mM imidazole. SmSTK25 (513-622) was buffer-exchanged into 1X PBS and 10% glycerol and applied to a Superdex 200 column for gel filtration chromatography on an AKTA FPLC. The sample was processed at a flow rate of 0.9 ml/min in 1X PBS and 10% glycerol. Eluate was collected as 90 1-mL fractions on a Frac 900 fraction collector (Amersham Pharmacia) maintained at 4°C. Each fraction was assessed for protein and concentrated with an Amicon concentrator, 10-kDa cut-off (Millipore). Rabbit polyclonal antibodies were generated by Cocalico Biologicals, Inc.

### Evaluation of kinase activity

For kinase assays with radiolabeled ATP, STK25 or STK25K48R (1.7μM) were incubated alone or together with either TAO or TAOK57R (0.3μM) and 50μM ATP ([γ-^32^P]ATP, 6,000-9,000 cpm/pmol) in 10mM Tris, pH 8.0, and 10mM MgCl_2_ for 10 min at 30°C. Following gel electrophoresis and autoradiography, STK25 or STK25K48R(1.25μM) as well as TAO or TAOK57R (0.25μM) bands were excised and analyzed by scintillation counting (Perkin Elmer, Tri-Carb 2910TR). For evaluation of STK25 phosphorylation by western blotting, proteins were incubated for 30 mins at 30°C in kinase assay buffer (10 mM Tris pH 8.0, 10 mM MgCl_2_, 0.5 μM per protein) with or without 50 μM ATP in a reaction volume of 30 μL. Reactions were quenched with 10μL of 4x Laemmli buffer and samples boiled at 99°C for 4 min, then stored at −20°C. Proteins were resolved by SDS-PAGE (Bio-Rad 4-20% precast polyacrylamide gel, cat# 4568095) for 45 min at 140V. The gel was placed in cold transfer buffer (25 mM Tris, 192 mM Glycine, 10% (v/v) methanol, pH ~8.4) and transferred to a nitrocellulose membrane (Bio-Rad cat# 1620115) for 60 min at 100V, 4°C. The membrane was stained with Ponceau S Solution (Sigma cat# P7170) for 5 min, imaged and destained by 2x washes with TBST (20 mM Tris, 150 mM NaCl, 0.1% (v/v) Tween20). The membrane was blocked for 1hr at RT with blocking buffer (Odyssey Blocking Buffer, Li-Cor cat# 927-40000) diluted 1:5 in TBS (20 mM Tris, 150 mM NaCl)), then stained O/N at 4°C with primary antibody diluted in blocking buffer. Membrane was washed 3x 5 min with TBST, then stained with secondary antibody diluted in blocking buffer, 1hr at RT. Membrane was washed 3x 5 min with TBST, then imaged with a LI-COR Odyssey imaging system. Primary antibodies were as follows: to detect phosphorylated T173 of smSTK25 and kinase-dead smSTK25, we used Rabbit-anti-MST4 + MST3 + STK25 (phospho T174 + T178 + T190) antibody [EP2123Y] (ab76579) from Abcam. To detect total smSTK25 and kinase-dead smSTK25 we used the Rabbit polyclonal antibody against the STK25 C-terminus described above. Secondary antibody for all blots was LI-COR IRDYE 680 red, Goat anti-Rabbit cat# 925-68071, and was used at a dilution of 1:10,000.

For mass spectrometry analyses of SmTAO phosphorylation of kinase-dead SmSTK25, kinase reactions were performed as above, and the SmSTK protein was excised from an SDS-PAGE gel and the protein was analyzed by the UT Southwestern Proteomics Core. Specifically, protein gel pieces were digested overnight with trypsin (Pierce) following reduction and alkylation with DTT and iodoacetamide (Sigma–Aldrich). The samples then underwent solid-phase extraction cleanup with Oasis HLB plates (Waters) and the resulting samples were analyzed by LC/MS/MS, using an Orbitrap Fusion Lumos mass spectrometer (Thermo Electron) coupled to an Ultimate 3000 RSLC-Nano liquid chromatography systems (Dionex). Samples were injected onto a 75 μm i.d., 75-cm long EasySpray column (Thermo), and eluted with a gradient from 1-28% buffer B over 90 min. Buffer A contained 2% (v/v) ACN and 0.1% formic acid in water, and buffer B contained 80% (v/v) ACN, 10% (v/v) trifluoroethanol, and 0.1% formic acid in water. The mass spectrometer operated in positive ion mode with a source voltage of 1.8 kV and an ion transfer tube temperature of 275 °C. MS scans were acquired at 120,000 resolution in the Orbitrap and up to 10 MS/MS spectra were obtained in the ion trap for each full spectrum acquired using higher-energy collisional dissociation (HCD) for ions with charges 2-7. Dynamic exclusion was set for 25 s after an ion was selected for fragmentation. Raw MS data files were converted to a peak list format and analyzed using the central proteomics facilities pipeline (CPFP), version 2.0.3 ^46,47^. Peptide identification was performed using the X!Tandem^48^ and open MS search algorithm (OMSSA) ^49^ search engines against the human protein database from Uniprot, with common contaminants and reversed decoy sequences appended ^50^. Fragment and precursor tolerances of 10 ppm and 0.6 Da were specified, and three missed cleavages were allowed. Carbamidomethylation of Cys was set as a fixed modification with oxidation of Met and phosphorylation of Ser, Thr, and Tyr set as variable modifications. Phosphorylation sites were localized using the ModLS algorithm, using cutoff values for positive site identification that represent a scenario where the false discovery rate is < 1% ^51^.

### Gene Ontology (GO)

The Gene Ontology (GO) annotation for *Schistosoma mansoni* was obtained from GeneDB (https://www.genedb.org/). GO term enrichment was performed using the weight01 method provided in topGO (v2.34.0) for biological process (BP), molecular function (MF) and cellular component (CC). For each category, the analysis was restricted to terms with a node size of >= 5. Fisher’s exact test was applied to assess the significance of overrepresented terms compared with the screened genes. The threshold was set as FDR < 0.05.

### Identification of *S. mansoni*-specific phenotypes

For data in **Supplementary Table 5**, orthologs of *S. mansoni* genes in *C. elegans*, *D. melanogaster* and human were identified from WormBase ParaSite^52^. We considered *S. mansoni* and *Schmidtea mediterranea* genes (taking the dd_Smed_v6 gene set from PlanMine^53^) to be one-to-one orthologs if they were each other’s top BLASTP hits, with *E*-value < 0.05, and the BLAST *E*-value of the top BLASTP hit was 105 times lower than the BLAST *E*-value for the next best hit. *C. elegans* RNAi/mutant phenotypes were identified from WormBase^54^ and *D. melanogaster* phenotypes from FlyBase^55^.

