## Supplementary Figures for "Large-scale RNAi screening uncovers new therapeutic targets in the human parasite *Schistosoma mansoni*"

### Extended Data Fig. 1

>Smp_002160
CTTCACGGTACCTAGGGGTTATTTTCACCTTTCTTTGCATTGTTTTTCAAACAATGGGTACTGCAATGGATGTCCTAGATATTCTCGATCTGAAGAAT-
CAAGTCCCAGAAAGTCTATGCTTGATAAAGAAGCTATTCTGAGTCGTGCAGAGAAAAGAAAGAGTTATCGTCCTAACCCACCACCAAAAAGGC-
CCGGTCATGTTCCTCGTGAAGTTTGGGGCTTACATAGCACTCTTAACAATAATCTTCCTCCGATTATGCCAACTGATAATACACCTCTTTATAAA-
C**AACCCAAAGCTGTGATTGGT**GTAGGACGAGTTCGTTCTTGGCAATGGACACCTTTTACAAATTCCGCTAGACAAGATAATCTAGTCTTATAT-
CATTGGAGACGAGAAAGCACTGATCCAGAGGCTAATAAAGATTATTATTTTGCACGATACAATAAGCATGTCACTGTTCCTGAATACACTGTAGAA-
GAGTACGAAACAATGCTAAAAGATCCAAAATGGAGTGAAGAGCGTACAGCACATTTGATGGAATTGGCTAAACGTTTTGATTTACGTTTTATTCATAT-
GAGAGATCGATGGGATTGTGAAAAGTTCCCTGGACGTCCATCGGTTGAAGATCTTAAAGAAAGATATTATGGAATTTTAACACAGTTAGATA-
AGGCTCGTGGAACAAATCTTTCACAAGGTCTAAGATATGATGCAGCACATGAACGTCGACGTAAACAACAATTAAGTTTACTTTATGGTCGTA-
CAAAAGATCAAGTTGAAGAAGAACAACGTTTAATTATGGAATTACGTAAAATCGAAGCTAGACGTAAAGAACGTGAACGTAAAAAACAAGATTTA-
CAAAAACTTATCTCATTGGCTGACAGTGTAACTCGGGATTGTGAATTATCTGAGCATACAGGTTATGAAACAAATACAGATGATACAAATTCTTG-
GCTTACA**TCTACACATTCGTCCGTTGG**CTGTGGTCTATCATCTTCCAATAATTCATTGTTAACATCAACAACACTAGCAGGAAATACTGGGTTA-
CAACGTAAGCGACCCGGTGGTGTTAGTGGTTTGAATGCTAGTAATAATAGTACTGGCAATATTACGTCATCGTCTAGTCAGTTAATTGGAAATA-
CAAATTCACTCATCAATACAGATTCTAGTTGTACGAGTTTAACCAGTACAGGAGCAGCTGGTAGCGCCAATACTGGAGCGGTGATATCCTCTG-
GAGATATTTGTGCTTCATTAAATTCTGCTTCGAATACAGCTAATACCGCATTGAATACACAATATTCAATAAACTTTTTGGATGTTAGCA**AAAATC-**
**CCGGTGCTCATCTA**CGTAGTCAAAAGATGAAATTACCCAGTAATTTAGGACAGAAAAAGACAAGAATCATTGAAAACTTCTTAGGTCATTTA-
CAAATTGATCCTAATCCACCGGCAACTGCAGAAATTGTTGAAGCATATAATGGTTTACGTTCAAAAATTCTACTTTTATTCGATTTACGTCTAGC-
GTTATTAAATTGTGACTTTGAATTACATTCAGCTCGACTAAGACTAGAAACATTTGCTCCTAATAAACCGTTACCTTGTGGACTTGCTAGTCTGC-
CAAATCCTAGTGTATCAGAATCACCTGAAGATTCTTATTATAATTTAATCGATCCGTTGATTTTGTCGGCACTTCGTGTAGCCGACTCAAAACGTC-
CAATTCTACCGCGTCATTCACTTGGAGTTGCTGGTACATTGGCTGCGGCTGCTGGAAGTTTAGGTTTGCATAATATGACTAGTGGAAGTAGT-
GGTATTCCATCTACTAATACAACAGTGGAGAAATCAGATCCAAAACAATGTCGATTGAACCCATCGTCCATTGTTAATTCTTCAGATAGTGGA-
CAGTATAATTCTAGTGGATTGACAACTACTCGATCGTCTCCAGCGTTGTCAGGAGGCGGAGGCACTGGGAATCTAGATTCAGATAATTCGTCAT-
CAGCCGGAGATGG**TATCGGCAGTTCACAACGTC**GTAGAAGAGCAGCAGCTTTAGAACAGGGACGTGTTTTAAAGAAACTTAAAATTAAAGGA-
CAACTTGTTGATTGAAACTGTTGGTTGTGGAAAAACAAACAAATTTTTTGTTGTCTATGTGTACATAAAAATATTTTTTTGATTGATCTCATTTGATC-
TACCGTTTCAGAAGGAAACAGTTTCATAATAAATAGTTTTCCCTC

Initial primers

T7 polymerase binding sequence | *NotI* | Gene-specific primer

Forward: GAATTTAATACGACTCACTATAGGGCGGCCGCAAAATCCCGGTGCTCATCTA

Reverse: GAATTTAATACGACTCACTATAGGGCGCGCCGACGTTGTGAACTGCCGATA

T7 polymerase binding sequence | *AscI* | Gene-specific primer

Primers for validation

Forward: AACCCAAAGCTGTGATTGGT

Reverse: CCAACGGACGAATGTGTAGA

**Extended Data Fig. 1. An example of strategy for a single gene from the initial RNAi screen and the subsequent validation.** PCR primers for the initial screening were modified by adding sequences including restriction enzyme sites (*NotI* on forward primer, *AscI* on reverse primer) and a T7 polymerase promoter on both forward and reverse primers. To validate PCR product sequences we digested with *NotI* and the amplicon was sequenced with a T7 primer. To validate the initial screening results, a new pair of primers for the target gene were designed to amplify a fragment that shows no overlap with the initial PCR product (magenta).

### Extended Data Fig. 2

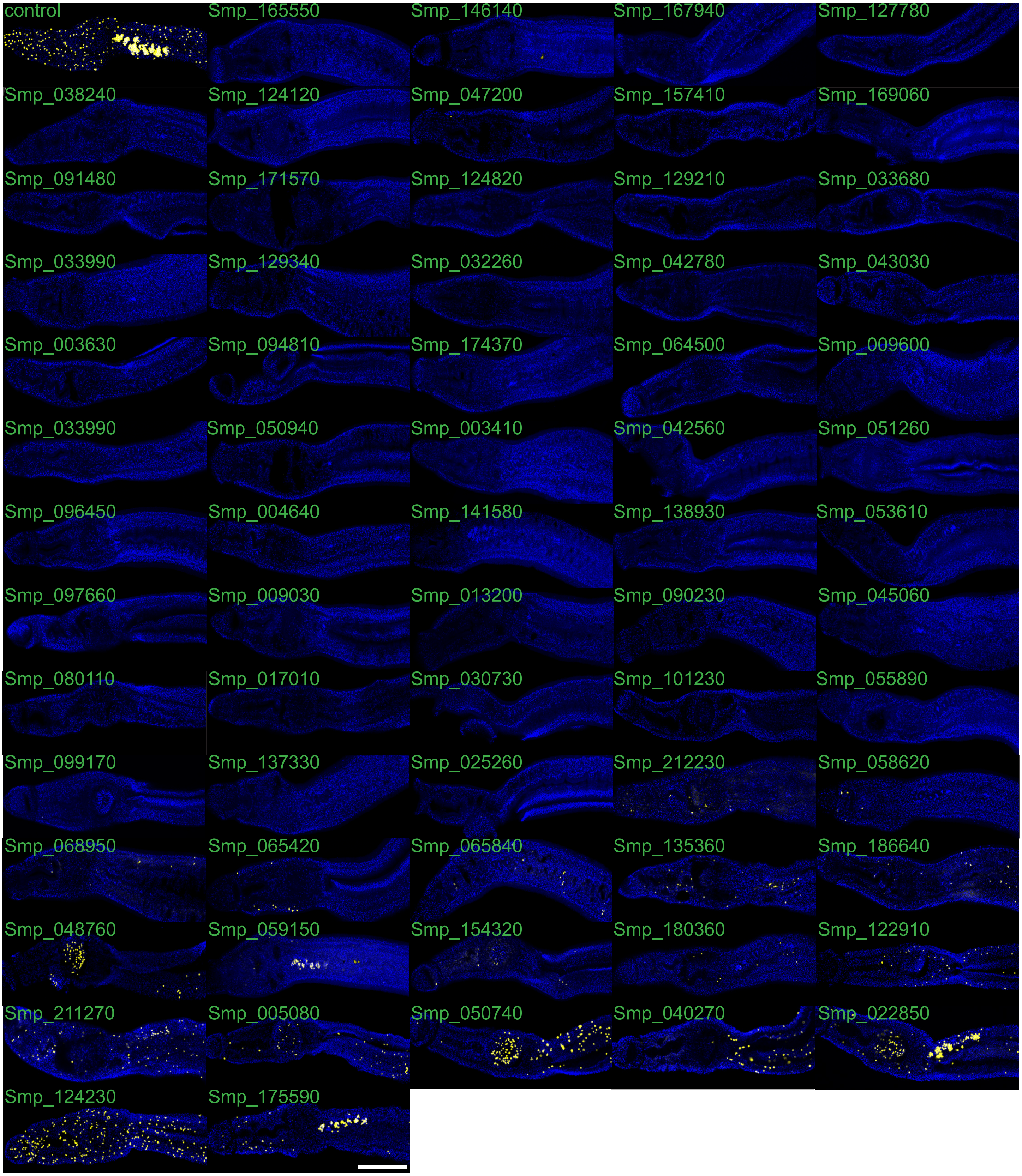


**Extended Data Fig. 2. All 66 hits with phenotypes of loss of stem cell maintenance.** EdU labeling of proliferative cells is shown in yellow, parasites are counterstained with DAPI to mark nuclei. Scale Bar, 200 µm.

### Extended Data Fig.3

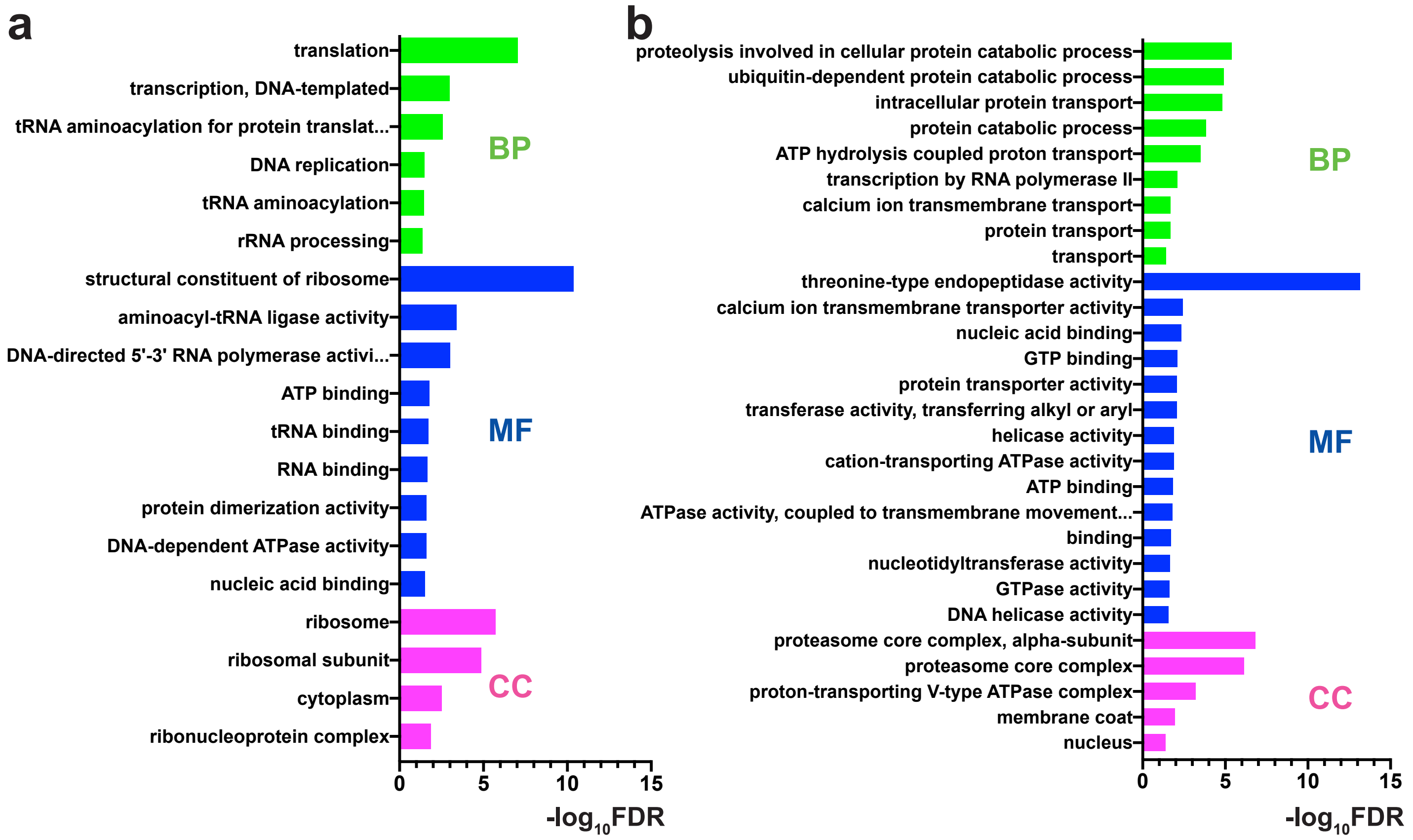


**Extended Data Fig. 3. Gene Ontology (GO) analysis. a**, GO analysis of all the 66 genes with a loss of stem cells phenotype. **b**, GO analysis of all the 195 genes with a detachment phenotype. GO term enrichment was performed by comparing hits with the entire list of 2,216 screened genes. All enriched GO terms with the threshold FDR< 0.05 are listed for Biological Process (BP), Molecular Function (MF) and Cellular Component (CC).

### Extended Data Fig. 4

**a**

**Ubiquitin/Proteasome System (UPS)**

**Ubiquitin system**

| Protein ubiquitination | Protein deubiquitination | Others |
|---|---|---|
| Smp_162630: E1 | Smp_069960: DUBs | Smp_018240: p97 |
| Smp_103710: E2 | Smp_089180: DUBs | |

**26S proteasome**

| 20S subunit | | 19S subunit |
|---|---|---|
| Smp_212920: α1 | Smp_074500: β2 | Smp_012470: Rpt1 |
| Smp_067890: α2 | Smp_121430: β3 | Smp_175250: Rpn2 |
| Smp_092280: α3 | Smp_056500: β4 | Smp_042270: Rpn3 |
| Smp_070930: α4 | Smp_212880: β5 | Smp_178810: Rpn9 |
| Smp_032580: α5 | Smp_034490: β6 | Smp_173840: Rpn10 |
| Smp_207080: α6 | Smp_073410: β7 | |
| Smp_076230: α7 | | |

**b**

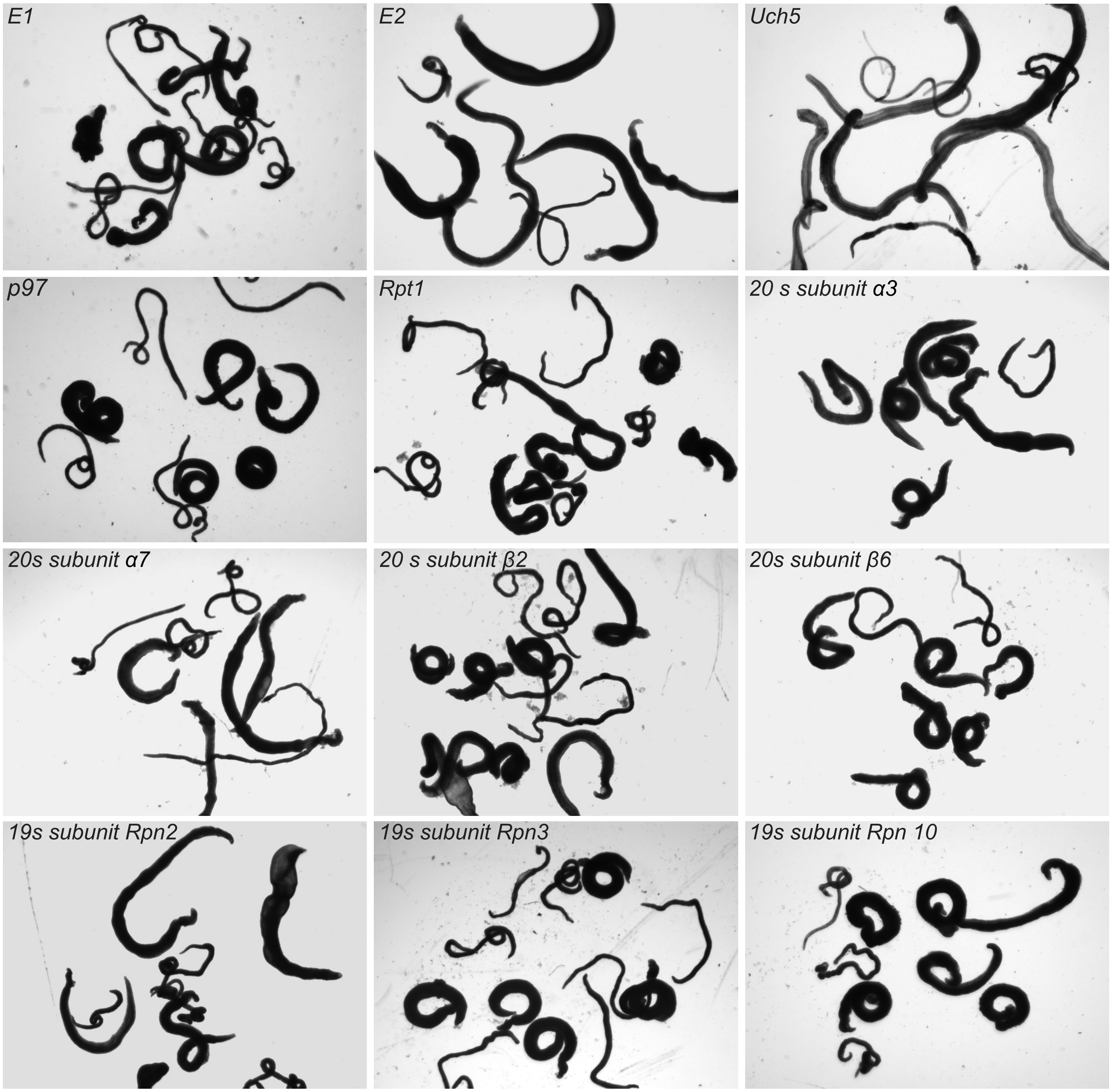


**Extended Data Fig. 4. Phenotypes of RNAi-treatment for various ubiquitin-proteasome system (UPS) components. a,** list of genes associated with visible phenotypes and the component of the UPS they encode**. b,** Knock down of listed UPS components in our 30-day treatment regime resulted in degeneration of the parasite's tegument or detachment from the substrate. *Uch5* (Smp_069960), ubiquitin carboxyl-terminal hydrolase 5, belongs to the deubiquitinaiting enzymes (DUBs).

### Extended Data Fig. 5

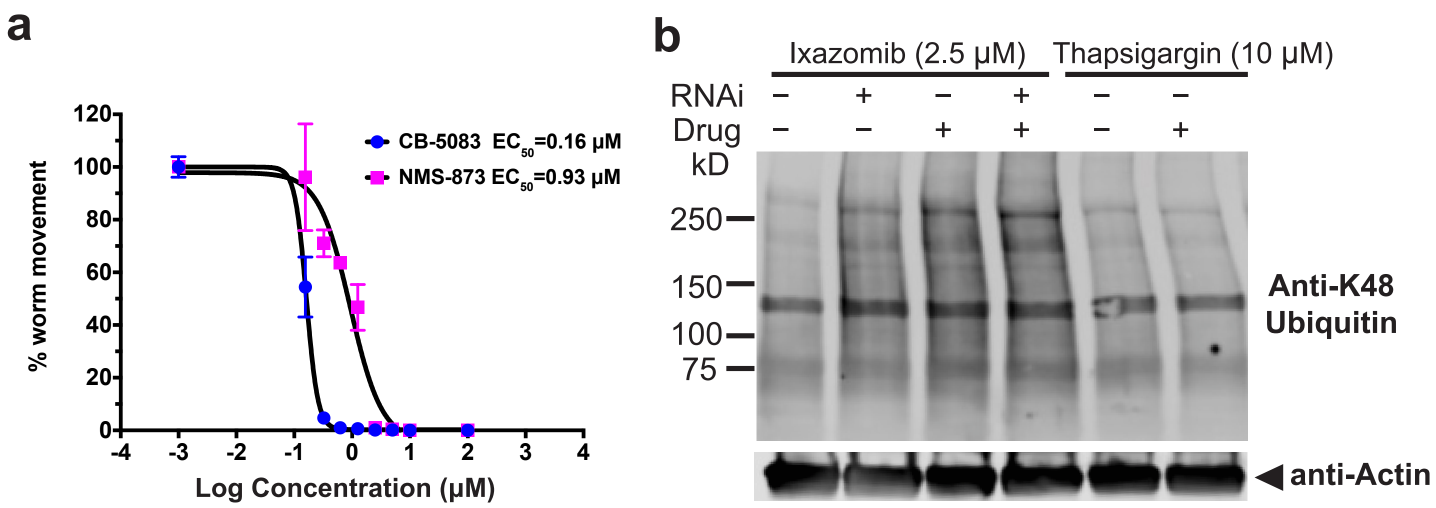


**Extended Data Fig. 5. Evaluation of p97/proteasome inhibitors. a,** EC50 values for the p97 inhibitors CB-5083 and NMS-873 on parasite movement. Error bars represent standard deviation of the mean motility scores (as percentages). n= 6 (three biological replicates, single adult worm pair/ well for each concentration point). **b,**Western Blot for K48-linked polyubiquitinated proteins. RNAi of *S. mansoni proteasome subunit beta type-2* or treatment of worms with the proteasome inhibitor Ixazomib caused an increase in the accumulation of K48-polyubiquitinated proteins. Treatment of worms with non-UPS inhibitor thapsigargin, which also compromised worm vitality, did not increase the polyubiquitinated proteins. Representative image from 3 replicates.

### Extended Data Fig. 6

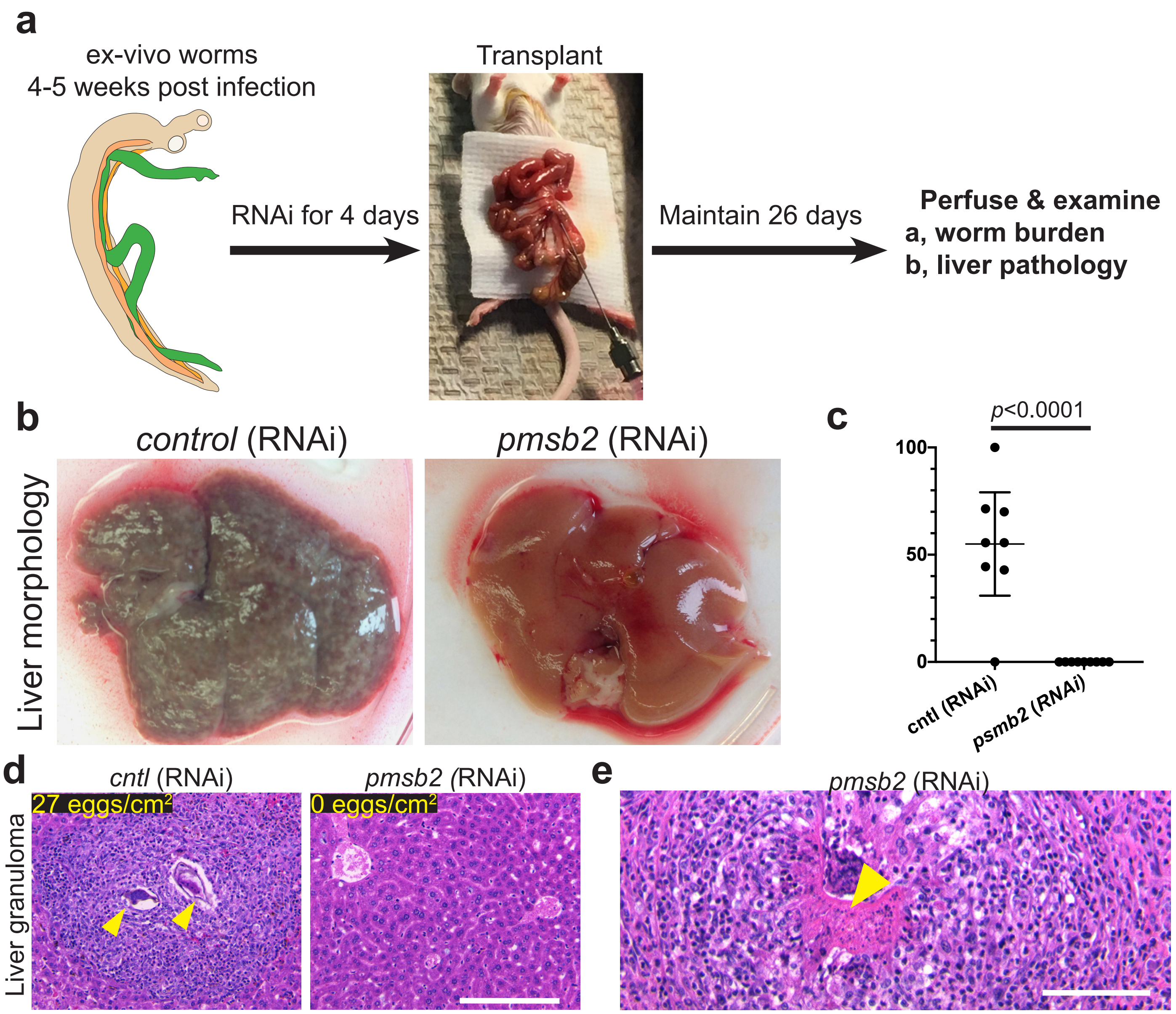


**Extended Data Fig. 6. *In vivo* validation of RNAi hits by surgical transplantation. a**, Strategy for *in vivo* validation. Parasites were perfused from mice at 4-5 weeks after infection and then treated with 30 μg/mL dsRNA for 4 days before surgical transfer into the mesenteric veins of naive mice. After 26 days, mice were sacrificed to harvest worms and livers. **b**, Liver morphology of mice after transplantation of worms that were pretreated with control dsRNA, *proteasome subunit beta type-2 (pmsb2)*. Granulomas were only observed in mice receiving control parasites. **c**, Percent recovery of male parasites treated with dsRNA specific to proteasome subunit beta type-2 (pmsb2, Smp_074500; n = 10 transplants) or an irrelevant dsRNA (control; n = 8 transplants) following surgical transplantation of parasites into mice. ****, $p < 0.0001$, t-test. **d**, Hematoxylin and Eosin staining of livers from recipient mice that received either control or pmsb2(RNAi). Schistosome egg-induced granulomas in livers were observed in control RNAi recipient mice, but not in pmsb2(RNAi) recipient mice. Counts of eggs per liver section are shown in top left, n=3.**e**, Transplanted parasites from proteasome *pmsb2*(RNAi) treatments were found trapped and in various stages of degeneration in livers of recipient mice. Scale bars, 100 μm.

### Extended Data Fig. 7

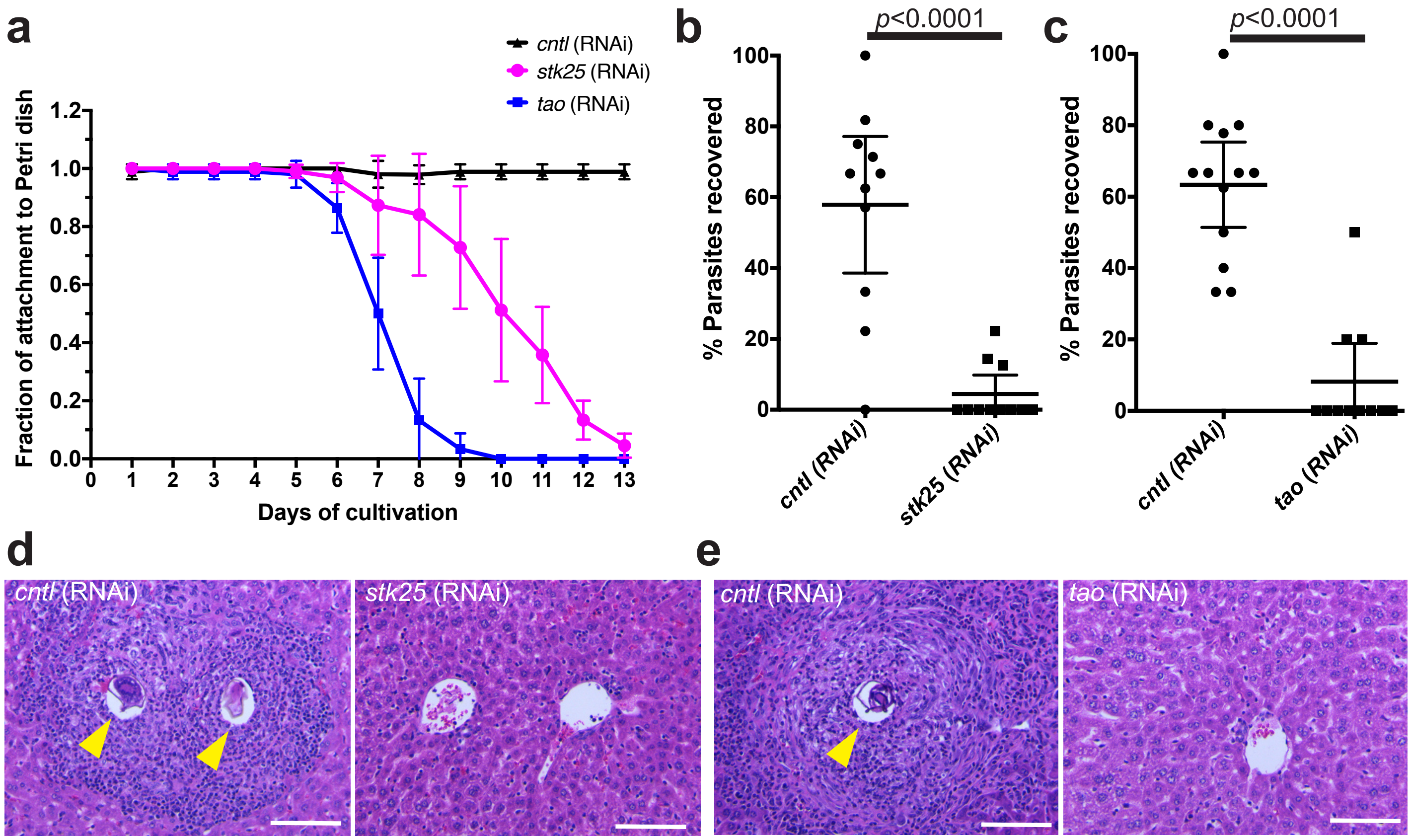


**Extended Data Fig. 7. Attachment curve of *stk25/tao* RNAi parasites *in vitro* and surgical transplantation of *stk25/tao* RNAi parasites. a,** dsRNA treatment of *S. mansoni* targeting either *stk25* or *tao* caused the worms to detach from Petri dish starting at 6 days after *in vitro* cultivation. cnlt, control. n=9 for each group with 3 replicates. Error bars represent 95% confidence intervals. **b,c,** Percent recovery of male parasites treated with dsRNA specific to *stk25* (Smp_096640; n=11 transplants) or *tao* (Smp_068060, n=11 transplants) following surgical transplantation of parasites into mice. For controls, n=11 transplants in **b** and n=13 transplants in **c**. ****, $p < 0.0001$, t-test. Error bars represent 95% confidence intervals. **d,e,** Hematoxylin and Eosin staining of livers from recipient mice that received either *cntl* or *stk25* (RNAi) or *tao* (RNAi). Egg-induced granulomas (indicated by yellow arrow) were only observed in control group. Scale bars, 100 µm.

### Extended Data Fig. 8

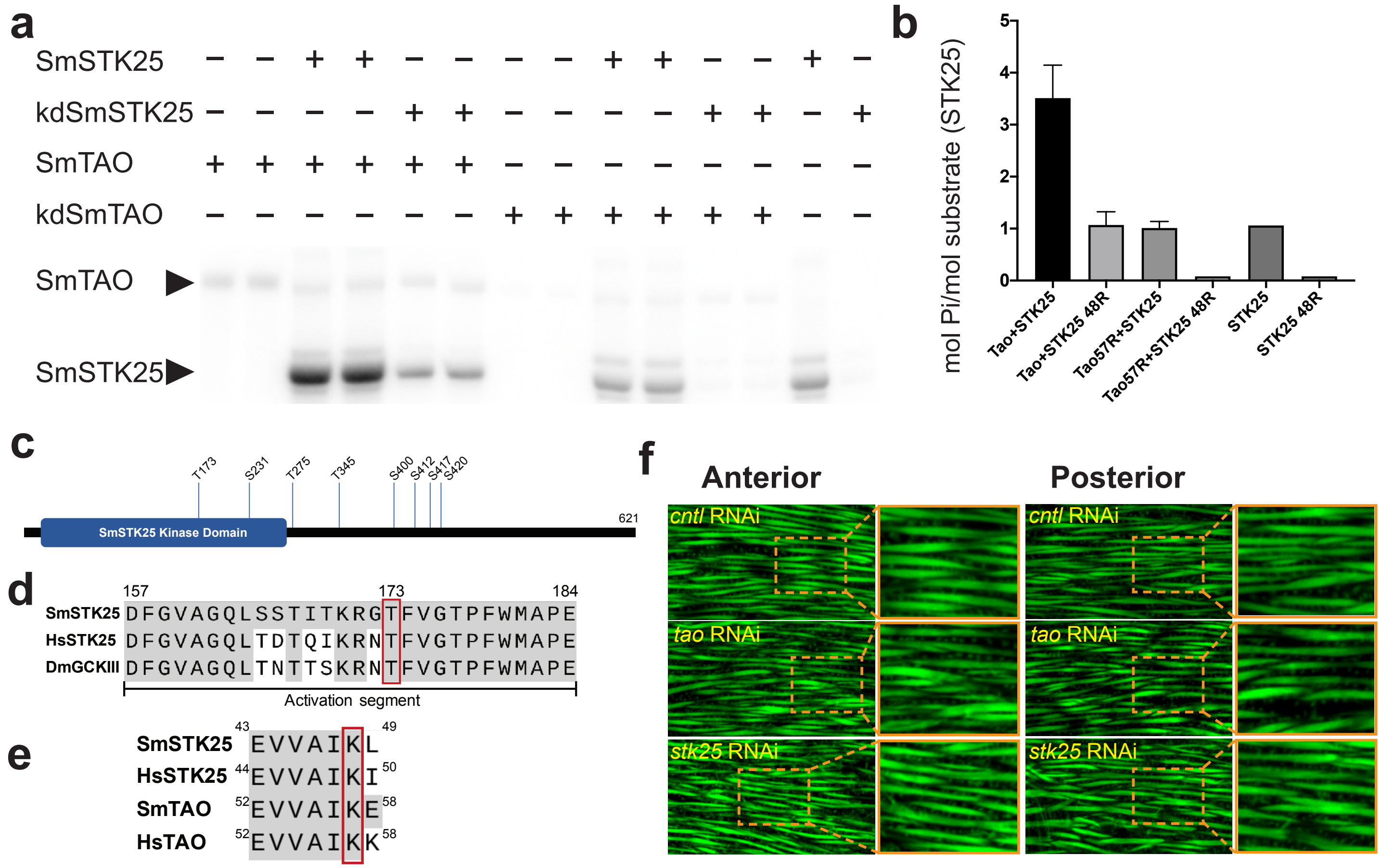


**Extended Data Fig. 8. Assays of *Schistosoma mansoni* protein kinases STK25 and TAO. a**, Autoradiography of kinase assay with recombinant schistosome proteins. Wild-type SmTAO protein phosphorylates both wild-type and kinase-dead SmSTK25, and that signal is ablated when wild-type SmTAO is replaced by a kinase-dead mutant (kdSmTAO, K57R). **b,** Quantification of radioactive phosphate transfer in kinase assay shown in (**a**), by scintillation counting. Error bars represent 95% confidence intervals. **c,** High confidence phosphosites of SmSTK25 kinase from mass spectrometry. To determine high confidence sites, only those with ModLS scores >30 and ID prob >0.95 on the PTM report were chosen. Eight high confidence phosphosites were detected and shown on the cartoon. **d,** Sequence alignment of *S.mansoni* STK25 kinase SmSTK25 activation segment and its human homologue HsSTK25 as well as *Drosophila* homologue DmGCKIII. Threonine 173 (T173) indicated in red box is the phosphosite in the activation segment highly conseved in different organisms. The positions of amino acids on SmSTK25 were used. **e**, Alignment of amino acids surrounding the VAIK motif required for catalysis in most kinases. To make kinase dead versions of SmSTK25 and SmTAO, the catalytic lysine in the VAIK motif was mutated to arginine (K48R and K57R for SmSTK25 and SmTAO, respectively). Sm, *Schistosoma mansoni*; Hs, *Homo sapiens.* **f**,Phalloidin staining of male parasites after dsRNA treatment with either Control (cntl) or *tao* or *stk25*. No obvious differences in muscle fiber morphology or numbers by phalloidin labeling were observed between *cntl* RNAi or *tao/stk*25 RNAi worms in either anterior (head) or posterior (body) parts of the worm body. Scale bars, 20 μm.

### Extended Data Fig. 9

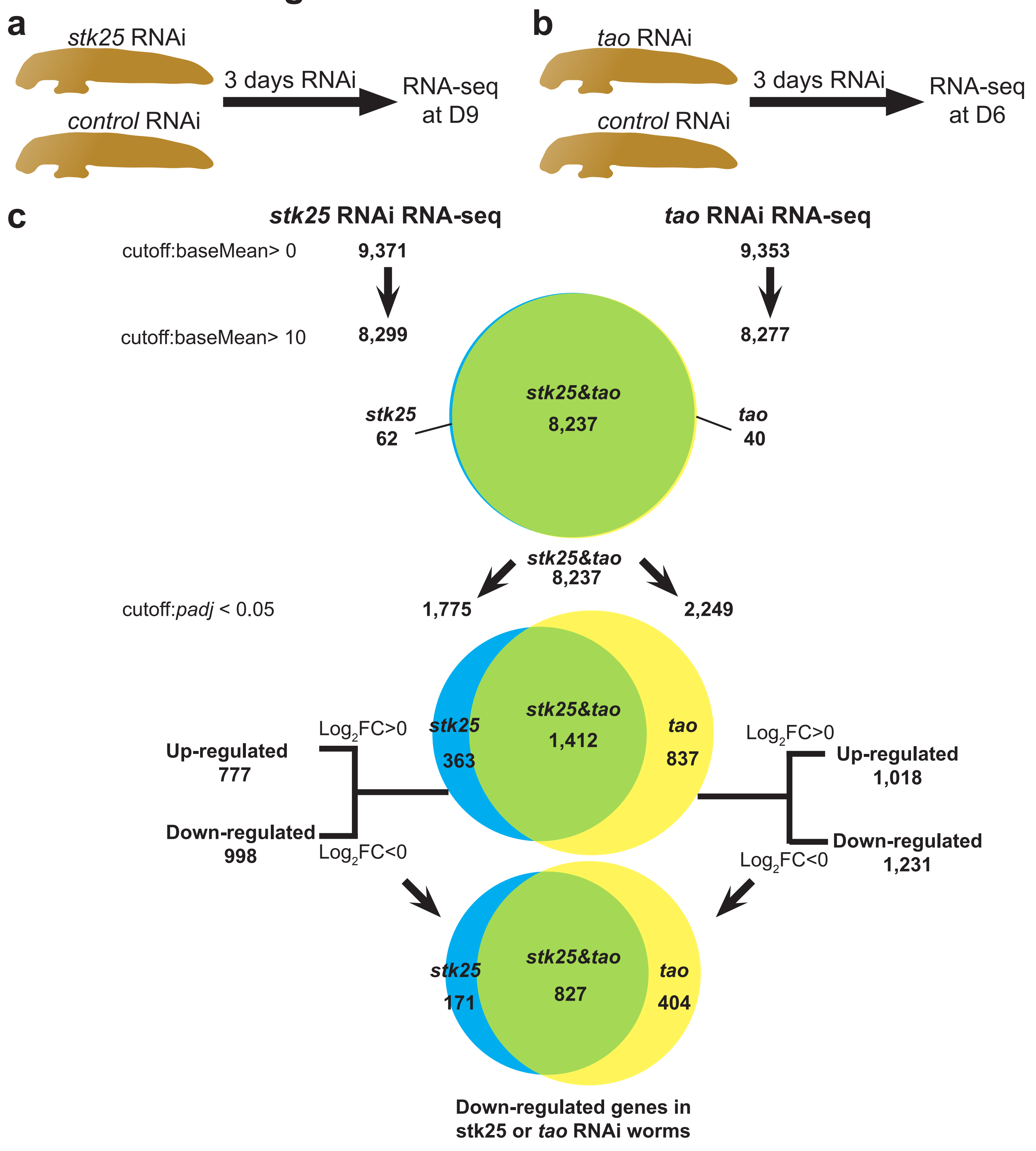


**Extended Data Fig. 9.RNA-seq and data analysis of the differentially expressed genes in *stk25/tao* RNAi**. **a**,**b**,RNAseq strategy for male worms treated with dsRNA that target either *tao* or *stk25*. **c**, Analysis of the differentially expressed genes after *stk25* or *tao* RNAi. Genes with low expression level (baseMean≤ 10) were excluded before further analysis. Among the 8,237 genes that are both expressed in *stk25* and *tao* datasets indicated by Venn diagram, 1,775 genes and 2,249 genes were significantly differentially expressed (*padj* <0.05) in either *stk25* RNAi or *tao* RNAi compared with control RNAi worms. Of the 998 and 1,231 down-regulated genes ($Log_2FC$ <0) in *stk25* RNAi worms or *tao* RNAi worms, 827 were shared in both groups.

### Extended Data Fig. 10

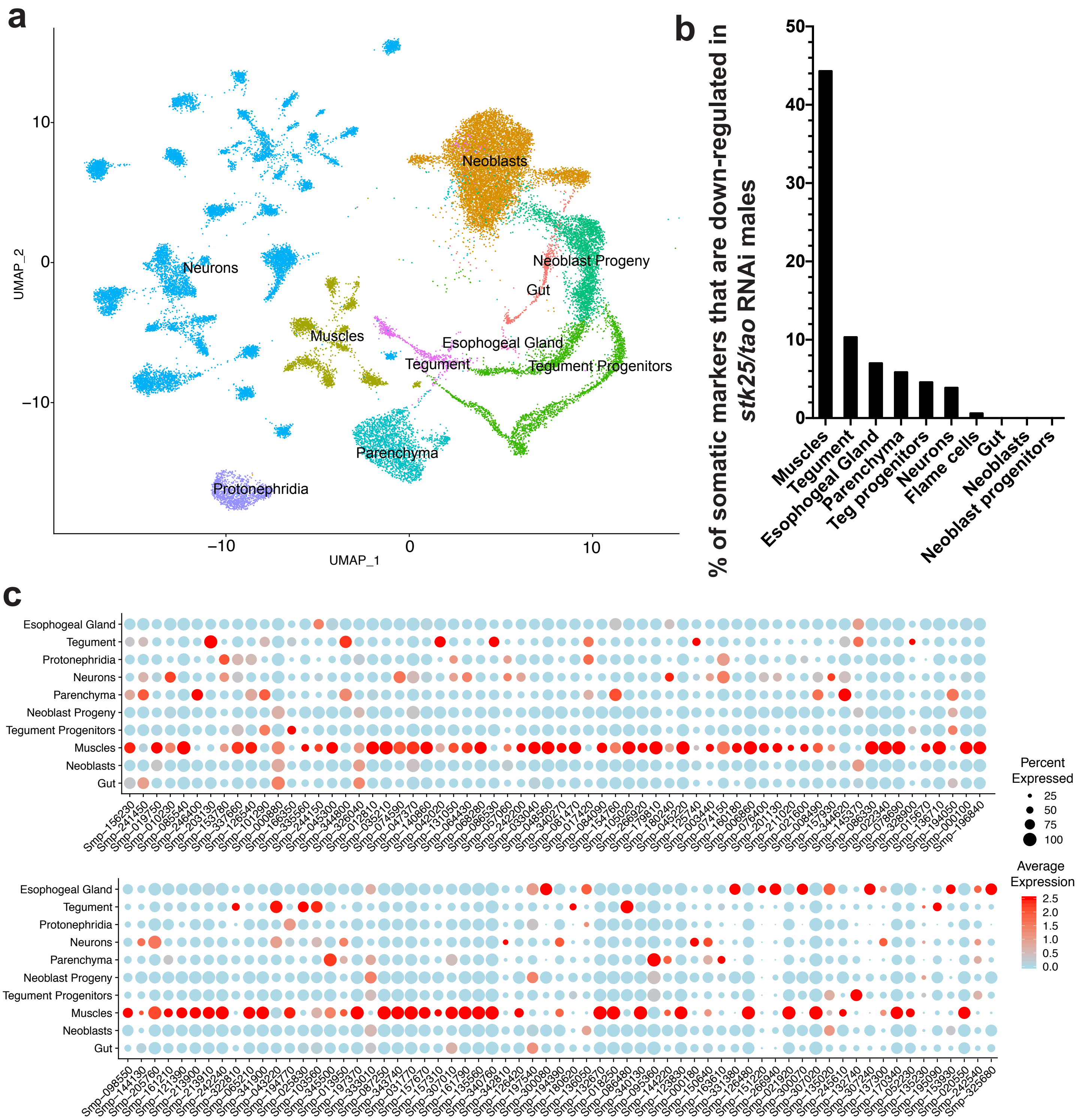


**Extended Data Fig. 10. Ratio of somatic markers in down-regulated genes in *stk25/tao* RNAi worms.** **a**, Dimensional reduction plot of male somatic tissue clusters from an adult single cell atlas. Each dot represents a cell and cells from the same cluster were labeled with same color. **b**, Percentage of tissue-specific somatic markers down-regulated (*padj* < 0.000001) after *stk25 or tao* RNAi treatment, measured by RNAseq. Nearly half of all muscle-specific markers are down-regulated in both *stk25 or tao* RNAi treatment groups. teg stands for tegument. **c**, Dotplot showing the expression of the top 129 down-regulated ($Log_2$ Fold Change <-0.5, adjusted *p*-value < 0.00001) genes in common to both *stk25* and *tao* RNAi datasets across somatic tissues. Most down-regulated genes are expressed in muscles.
